# A novel role of 3’,5’-cAMP in the regulation of actin cytoskeleton in Arabidopsis

**DOI:** 10.1101/2022.01.31.478439

**Authors:** Monika Chodasiewicz, Nicolás E. Figueroa, Peter Franz, Marcin Luzarowski, Juan C. Moreno, Dorothee Childs, Aleksandra Masiuk, Manny Saluja, Arun Sampathkumar, Georgios Tsiavaliaris, Aleksandra Skirycz

## Abstract

The role of cyclic adenosine monophosphate (3’,5’-cAMP) in plants is not well understood, and here, we report a novel role of 3’,5’-cAMP in regulating the actin cytoskeleton. The 3’,5’-cAMP treatment affects the thermal stability of 51 proteins, including a vegetative actin isoform, ACTIN2. Consistent with the above results, the increase in 3’,5’-cAMP levels, obtained either by feeding or by chemical modulation of 3’,5’-cAMP metabolism, is sufficient to partially rescue the short hypocotyl phenotype of the actin2 actin7 mutant, severely compromised in actin level. No such complementation was measured for a positional isomer of 3’,5’-cAMP, 2’,3’-cAMP, attesting to the specificity of 3’,5’-cAMP treatment. Moreover, supplementation of 3’,5’-cAMP partly counters the activity of an actin-depolymerizing drug latrunculin B. In vitro characterization of the 3’,5’-cAMP – actin interaction argues against the direct binding. Instead, based on the proteomics characterization of the *act2act7* mutant supplemented with 3’,5’-cAMP, we hypothesize that 3’,5’-cAMP affects cytoskeleton dynamic by modulation of calcium signaling, and actin binding proteins.

## Introduction

The second messenger adenosine 3’,5’-cyclic phosphate (3’,5’-cAMP) was first reported in mammalian cells in 1957 (Rall and Sutherland, 1958). Shortly thereafter, the signaling role of 3’,5’-cAMP was also described in both yeast (Sy and Richter, 1972) and bacteria (Makman and Sutherland, 1965). 3’,5’-cAMP is produced from ATP *via* activity of the membrane-localized adenylyl cyclases (AC) (Smith et al., 1977), and degraded into AMP by the enzyme phosphodiesterase (PDE) (Genschik et al., 1997). In animals, 3’,5’-cAMP works by activating protein kinase A (PKA) to enable the phosphorylation of a multitude of protein substrates involved in the regulation of cell growth, metabolism, and stress resistance (Turnham and Scott, 2016). Additional 3’,5’-cAMP targets include a family of cyclic nucleotide-gated cation channels (CNGCs) (Yau, 1994) and nucleotide exchange factors, named Epac (exchange factor directly activated by cAMP) (de Rooij et al., 1998). The integration of the EPAC- and PKA-dependent pathways governs the net physiological effects of 3’,5’-cAMP signaling (Cheng et al., 2008).

In contrast to animal and yeast cells, the function of 3’,5’-cAMP in plants remains enigmatic (Gehring and Turek, 2017). Although plants unequivocally produce 3’,5’-cAMP (Amrhein, 1977; Brown and Newton, 1981), albeit in nM (Moutinho et al., 2001) rather than the μM concentrations reported for other eukaryotes, bioinformatics searches have failed to identify plant homologs of animal PKA and Epac proteins (Gehring and Turek, 2017). Similarly, despite evidence supporting their activity, adenylate cyclase (AC) and phosphodiesterase (PDE) have not been isolated and/or fully characterized in plants (Bianchet et al., 2019; Gehring and Turek, 2017). To date, the only known and functionally validated plant 3’,5’-cAMP protein interactors are CNGCs (Duszyn et al., 2019; Jarratt-Barnham et al., 2021; Zelman et al., 2012). Binding of 3’,5’-cAMP to the intracellular part of the channel leads to an increase in channel conductance and thereby affects cellular ion homeostasis (Duszyn *et al*., 2019; Leng et al., 2002; Talke et al., 2003; Zelman *et al*., 2012). In fact, the first functional evidence for a specific regulatory role for 3’,5’-cAMP in plants came from experiments that demonstrated the 3’,5’-cAMP dose-dependent activation of a K^+^ channel activity in *Vicia faba* mesophyll protoplasts (Li et al., 1994). The characterized members of the CNGC family have been shown to conduct Na^+^, K^+^, and/or Ca^2+^ ions, and are involved in plant responses to both biotic and abiotic stresses. For instance, K^+^ and Na^+^ permeable CNGC4 (Balague et al., 2003) is important for building up the hypersensitive response associated with disease resistance, while CNGC2 (Lu et al., 2016) and CNGC6 (Gao et al., 2012) conduct Ca^2+^ ions in response to jasmonic acid and heat stimuli, respectively.

In the past and in the absence of the known metabolic enzymes or signaling components, the strategies used to study the role of 3’,5’-cAMP in plants have relied on the use of cell-permeable 3’,5’-cAMP analogs (Alqurashi et al., 2016; Jiang et al., 2005), and compounds such as forskolin (Totsuka et al., 1983) and theophylline (Dryden et al., 1988) that are recognized as positive regulators of 3’,5’-cAMP metabolism. More recently, and to overcome the limitations of using chemical compounds, an elegant approach based on the over-expression of so-called ‘cAMP-sponge’ (cAS) was developed in *Arabidopsis* (Sabetta et al., 2019) and tobacco cell cultures (Sabetta et al., 2016). The cAMP sponge is composed of the two high affinity cAMP binding domains (Lefkimmiatis et al., 2009) that effectively sequester the active pool of cellular 3’,5’-cAMP to yield transgenic lines defective in 3’,5’-cAMP signaling/regulation (Sabetta *et al*., 2016). Phenotypic and molecular analysis of the cAMP sponge lines supported previous work (Balague *et al*., 2003; Ehsan et al., 1998; Yoshioka et al., 2006) implicating importance of 3’,5’-cAMP signaling in the regulation of cell cycle progression and plant resistance against bacterial pathogens (Sabetta *et al*., 2019; Sabetta *et al*., 2016).

The identification of 3’,5-cAMP protein receptors, other than GNGCs, has daunted plant scientist for the last 20-years. In the past, and following the success of analogous approaches in animal cells (Aye et al., 2009), affinity purification (AP) using beads coupled to 3’,5’-cAMP was also conducted in plants (Donaldson et al., 2016). To date, the most comprehensive study yielded a list of 15 putative protein interactors, mainly metabolic enzymes, for future characterization (Donaldson *et al*., 2016). Here, we decided on an alternative biochemical approach that circumvents the need for compound immobilization, which can interfere with binding. Thermal proteome profiling (TPP) (Savitski et al., 2014) combines thermal stability shift assays and untargeted proteomics. Protein interactors are selected based on the change in their melting temperature upon ligand binding. TPP is known for its specificity, and TPP/3’,5’-cAMP experiments in mammalian cells have successfully retrieved known 3’,5’-cAMP interactors (Lim et al., 2018). Following on successful use of TPP for deciphering cAMP binding proteins in mammals, we used this TPP to characterize 3’,5’-cAMP interactome in *A. thaliana*. From the identified putative 3’,5’-cAMP interactors, we focused on actin. And however, our results argue against actin being a direct target of 3’,5’-cAMP; we could demonstrate that 3’,5’-cAMP does affect actin cytoskeleton, likely downstream of calcium signaling hence CNGCs. Our results clearly shows that 3’,5’-cAMP restore *act2act7* short hypocotyl phenotype, and that it can counteract Lat B effect on plant growth. Finally we also proved that modulation of endogenous level of 3’,5’-cAMP affects hypocotyl growth.

## Results

### 3’,5’-cAMP affects thermal stability of ACTIN2

Here, we used TPP (Savitski *et al*., 2014) to identify 3’,5’-cAMP protein interactors. TPP is a biochemical approach that enables monitoring of small molecule–protein interactions in a native protein lysate by exploiting the fact that each protein unfolds (‘melts’) and aggregates at a defined temperature, referred to as its melting temperature (T_m_). Typically, ligand binding affects the tertiary structure of the protein, thereby affecting the T_m_ in either a stabilizing or destabilizing manner. A TPP protocol, in brief, includes heating of a native lysate, with or without the studied ligand, to different temperatures, followed by short centrifugation (**Figure 1a**). Denatured proteins (the aggregates) migrate to the pellet, whereas proteins that retain their proper folded configuration remain in the soluble fraction. The proportion of a given protein remaining in the soluble fraction across the different temperatures is quantified and used to plot protein-melting curves. The temperature at which 50% of the protein is unfolded is defined as T_m_.

**Figure 1.**
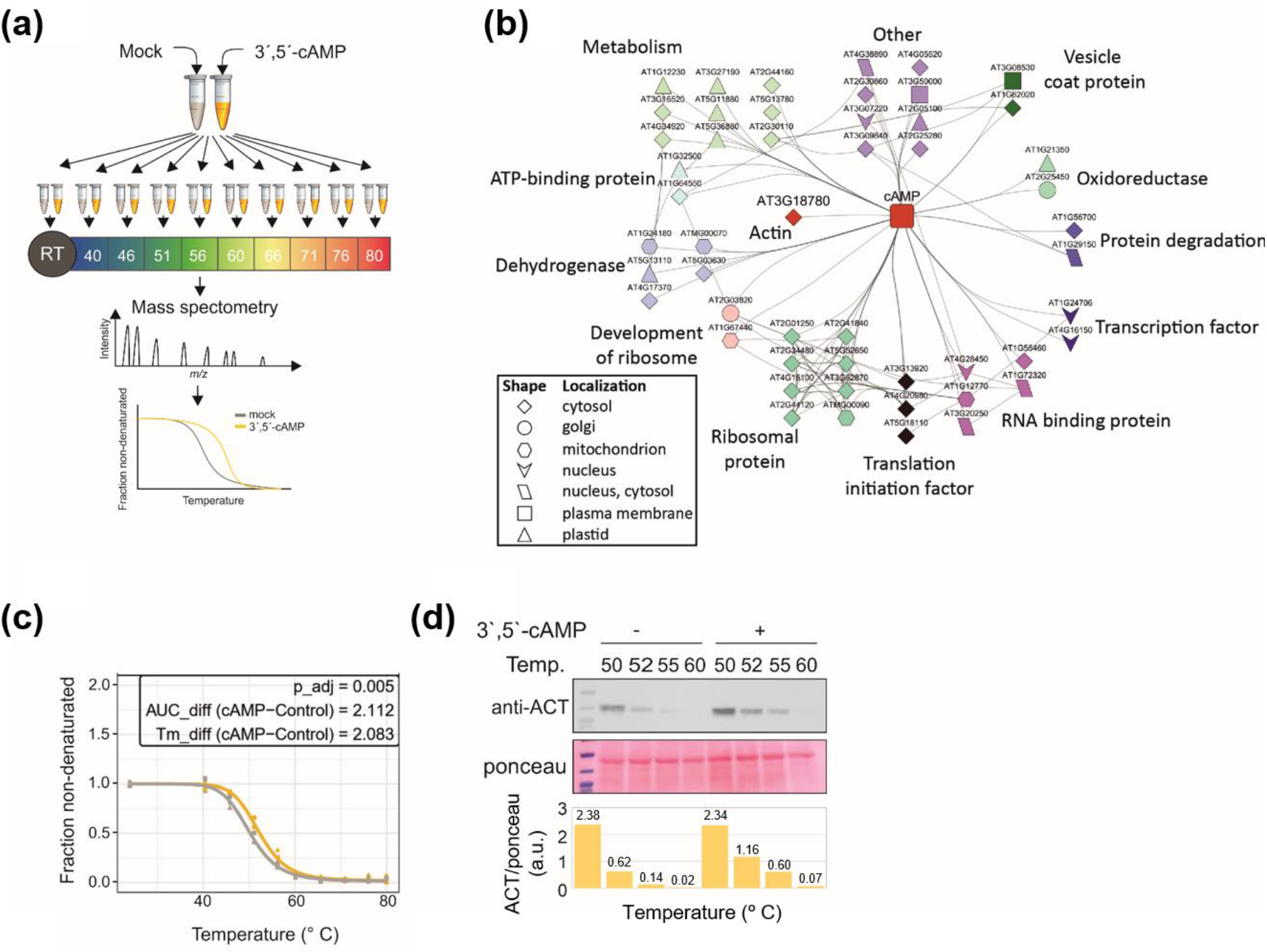
Thermal proteome profiling experiment identified 51 putative 3’,5’-cAMP binding proteins. (**a**) Experimental design. (i) Cell lysate prepared from the Arabidopsis cell culture was first treated with 10 μM 3’,5’-cAMP (ligand) or DMSO (mock), (ii) equal aliquots were heated to different temperatures in a temperature gradient ranging from 40 °C to 80 °C (plus RT control), (iii) brief centrifugation was used to separate denatured from non-denatured (soluble) proteins, (iv) soluble proteins were precipitated, trypsin digested, and analyzed by mass spectrometry, (v) Melting curves for mock and ligand samples were fitted and used to calculate Tm. The figure was modified from (Franken et al., 2015). (**b**) Interaction network of the 51 putative 3’,5’-cAMP binders delineated using TPP. Information was retrieved from String (von Mering et al., 2003) using experimental, database and literature evidence (score > 0.4). Network was prepared in Cytoscape (Shannon et al., 2003). (**c**) Melting curve for ACTIN 2 (ACT2) in the presence (yellow line) and absence (grey line) of 3’,5’-cAMP. Data are from three replicates. Replicate corresponds to independent 3’,5-cAMP treatments but with the same cell lysate. (**d**) Western blot analysis of the actin stability in the TPP experiment, in the temperature gradient ranging from 50 °C to 60 °C. For the complete TPP dataset, see **Supplementary Tables 3 and 4**. Western blots were quantified using Image J. Data are available in **Supplementary Table 5**.

We chose Arabidopsis cell cultures as a source of starting material as 3’,5’-cAMP was associated with the regulation of cell cycle progression (Ehsan *et al*., 1998; Sabetta *et al*., 2016). We also used 10 μM 3’,5’-cAMP, which is approximately 50 times the reported cellular concentrations (Ashton and Polya, 1978), and ten different temperatures, starting from room temperature (RT) as a control and then using a step-gradient from 40°C to 80°C. Of the 4983 identified proteins (**Supplementary Table 3**), 4682 were quantified in all three replicates and treatment groups and were subjected to melting curve fitting, using the hypothesis testing described by (Childs et al., 2019). Of the 4295 proteins for which melting curve fits and F-test *p*-values could be determined, 51 showed significant changes in curve shape between the control and treatment group (adjusted F-test *p*-value ≤0.05) (**Figure 1b, Supplementary Table 4**). Of these, 46 proteins were stabilized and five were destabilized. The identified proteins belonged to different functional groups (**Figure 1b**), including multiple ribosomal subunits, and metabolic enzymes.

However, the protein that drew our attention was ACTIN 2 (ACT 2) as the regulatory connection between 3’,5’-cAMP and actin cytoskeleton has been reported before in animal cells e.g. (Gerits et al., 2007). 3’,5’-cAMP stabilized ACT2 by approximately 2°C (**Figure 1c**), and actin stabilization in the presence of 3’,5’-cAMP was validated in an independent experiment using western blotting and actin antibodies (**Figure 1d, Supplementary Table 5**).

### Supplementation with 3’,5’-cAMP rescues the short-hypocotyl phenotype of the actin *act2act7* mutant

The *Arabidopsis* genome encodes for eight *ACTIN* isoforms: *ACT2, ACT7*, and *ACT8* are vegetative, while the other five are reproductive (Gilliland et al., 2003; Kandasamy et al., 2009). *ACT2* and *ACT7* are highly abundant, whereas *ACT8* expression is relatively weak. A double mutant of *act2* and *act7* (*act2act7*) shows severely compromised actin abundance and is characterized by delayed development and dwarfed stature, with a short hypocotyl and only a few small leaves (Kandasamy et al., 2012).

As demonstrated above, treatment with 3’,5’-cAMP affected actin thermostability. We therefore wondered whether Br-3’,5’-cAMP supplementation could at least partially complement the *act2act7* growth phenotype by affecting actin dynamics. Br-3’,5’-cAMP is a membrane-permeable version of 3’,5’-cAMP (Alqurashi *et al*., 2016). To this end, we examined the hypocotyl length of dark-grown *act2act7* and WT seedlings grown on MS medium with and without 10 μM Br-3’,5’-cAMP. Br-3’,5’-cAMP supplementation significantly increased the hypocotyl length of the five-day-old *act2act7* seedlings, leading to the partial rescue of the short-hypocotyl phenotype. (**Figure 2a-d, Supplementary Figure 1, Supplementary Table 6**). Notably, Br-3’,5’-cAMP supplementation had no effect on the hypocotyl length of the WT plants (**Figure 2a,b, Supplementary Figure 1, Supplementary Table 6**). Next, we measured the hypocotyl length every day for up to five days after sowing (das) to determine when the growth difference appears.

**Figure 2.**
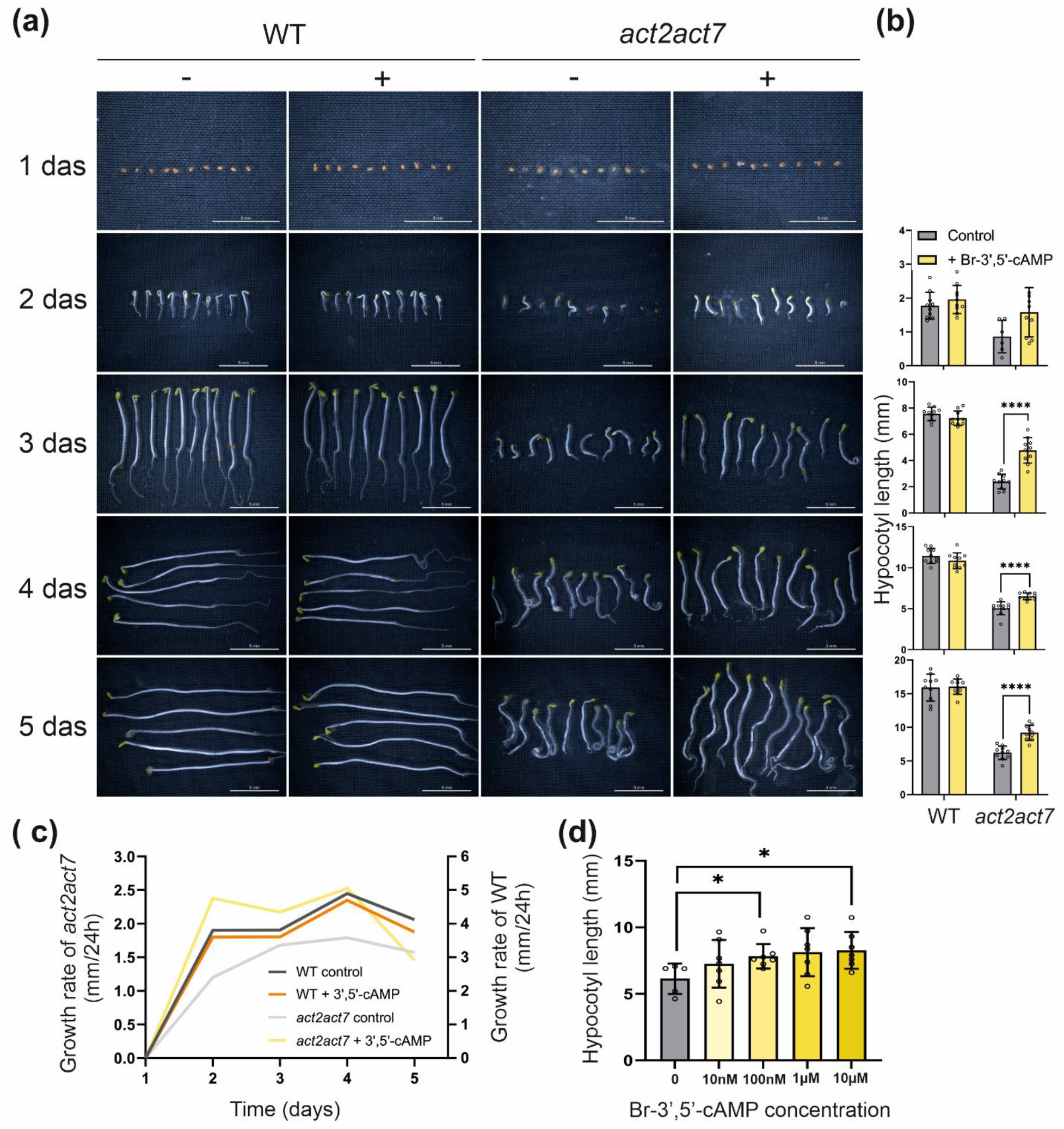
3’,5’-cAMP restores hypocotyl growth of the *act2act7* mutant. (**a**) WT and *act2act7* seedlings grown in the dark on control MS medium (-) or MS medium supplemented with 10 μM Br-3’,5’-cAMP (+). WT - wild type, *act2act7* - double *actin 2 actin 7* mutant. Scale bar = 5mm. (**b**) Hypocotyl length measurements at two to five days after sowing (das) (**Supplementary Table 6**). Hypocotyl length is expressed in mm. Values are given as means ± SD (n=10), except *act2act7* at day 2 (-) (n=5). Student’s t-test, * indicates p-values ≤ 0.05, and **** indicates p-values ≤ 0.0001. (**c**) Growth rate of WT and *act2act7* plants grown on control MS plates and plates supplemented with Br-3’,5’-cAMP. A rolling average was used to smooth the data. (**d**) Hypocotyl length measurements taken from five-day-old *act2act7* plants grown in the dark on control MS medium (0) or supplemented with different concentrations of Br-3’,5’-cAMP (**Supplementary Table 8**). Values are given as mean ±SD, n=5 for control, n=7 for treatment. Photographs can be found in **Supplementary Figure 3**. Asterisk indicates significant differences defined by Student’s T-test, *p*-value ≤0.05.

Calculation of the growth rate, defined as mm of growth per 24 h (mm/24h) revealed that treatment with 10 μM Br-3’,5’-cAMP had the most pronounced effect on the *act2act7* mutant early on, shortly after germination (**Figure 2C**). To exclude the possibility that 3’,5’-cAMP rescue is due to the 3’,5’-cAMP degradation to adenosine (Jackson, 1991) we grew *act2act7* seedlings on Br-adenosine. The treatment with 10 μM Br-adenosine (**Supplementary Figure 2**) did not rescue the short-hypocotyl phenotype of the *act2act7* mutant, nor did treatment with positional isomer of 3’, 5’-cAMP, 10 μM Br-2’,3’-cAMP (**Figure 3a,b; Supplementary Table 7**), asserting to the specificity of the 3’,5’-cAMP action. Finally, we showed that Br-3’,5’-cAMP concentrations as low as 100 nM, corresponding to the reported 3’,5’-cAMP concentrations (low to middle range nM) reported in plant cells (Moutinho *et al*., 2001), rescued the short-hypocotyl phenotype of the *act2act7* mutant (**Figure 2d**, **Supplementary Figure 3, Supplementary Table 8**).

**Figure 3.**
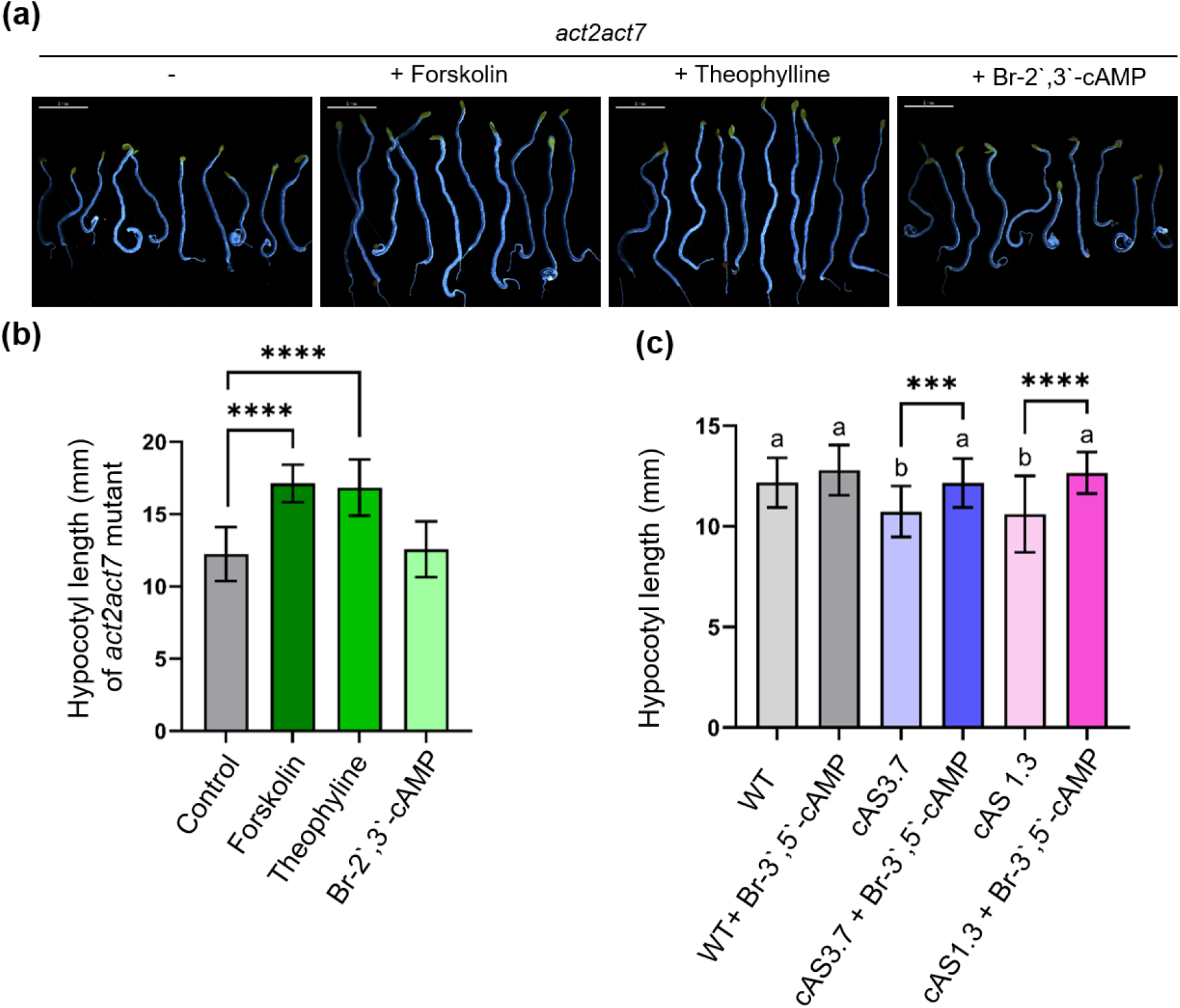
Manipulation of endogenous levels of 3’,5’-cAMP affects hypocotyl growth of *act2act7* and cAS lines. (**a**) *act2act7* seedlings grown in the dark on control MS medium (-) or MS medium supplemented with 10 μM forskolin, 100 μM theophylline or 10 μM Br-2’,3’-cAMP. Scale bar = 5mm. (**b**) Hypocotyl length measurements of *act2act7* seedlings growing for seven days in the dark (panel a)). Hypocotyl length is expressed in mm. Values are given as means ± SD; n=20, except for the treatment with Br-2’,3’-cAMP, where n=10. Student’s t-test, **** indicates p-values ≤0.0001. Measurements are included in **Supplementary Table 7**. Images of WT hypocotyls are included in **Supplementary Figure 4**. (**c**) Hypocotyl length of WT and sponge lines (cAS) growing in the dark for five days on control medium or supplemented with 10 μM Br-3’,5’-cAMP. Hypocotyl length is expressed in mm. Values are given as means ± SD; n=20 for WT and n= 32–36 for cAS lines. Letters and asterisks represent significant differences, Student’s t-test. *** indicates p-values ≤ 0.001, and **** indicates p-values ≤ 0.0001. Images of hypocotyls are included in **Supplementary Figure 5**, and measurements are included in **Supplementary Table 9**.

### Manipulation of the cellular 3’,5’-cAMP levels affects hypocotyl growth

To complement the Br-3’, 5’-cAMP experiments, we sought to understand how the endogenous 3’,5’-cAMP affects actin cytoskeleton and plant growth by examining hypocotyl lengths of the dark-grown *act2act7* mutant grown on plates supplemented with compounds known to increase the endogenous level of 3’,5’-cAMP. Forskolin acts by activating adenylyl cyclase, the enzyme responsible for 3’,5’-cAMP biosynthesis (Daly, 1984). Theophylline, an inhibitor of phosphodiesterases, boosts endogenous 3’,5’-cAMP levels by interfering with 3’,5’-cAMP degradation (Dryden *et al*., 1988). The *act2act7* and WT plants were grown in the dark on MS medium supplemented with either 10 μM forskolin or 100 μM theophylline. Seven-day-old seedlings were used for the hypocotyl length measurements. The elevation of endogenous 3’,5’-cAMP levels increased hypocotyl length in the *act2act7* mutant (by ~4-5 mm) but not in the WT plants (**Figure 3a,b; Supplementary Table 7, Supplementary Figure 4**).

The lack of effect of increases in the 3’,5’-cAMP levels on the hypocotyl length of the WT seedlings indicated that the endogenous 3’,5’-cAMP concentration is likely saturating. Therefore, we reduced the amount of 3’,5’-cAMP in the cell using Arabidopsis lines overexpressing the cAMP-sponge (cAS plants) (Sabetta *et al*., 2019). The cAS lines had a total 3’,5’-cAMP content similar to that of the WT plants; however, the level of free (and thus functionally relevant) 3’,5’-cAMP was reduced by 40–50% (Sabetta *et al*., 2019). Two different cAS lines (1.3 and 3.7) and a corresponding WT were grown for five days in the dark to examine whether the reduction of free 3’,5’-cAMP levels would affect hypocotyl length. Both cAS lines showed small, but significant, reductions of 1.5–2 mm in hypocotyl length in comparison to the WT (**Figure 3c, Supplementary Figure 5, Supplementary Table 9**). Supplementation with 10 μM Br-3’,5’-cAMP restored the short-hypocotyl phenotype of the cAS lines, confirming that the observed effect was due to 3’,5’-cAMP deficiency.

### 3’,5’-cAMP supplementation promotes actin bundling in Arabidopsis hypocotyl cells

In order to study the effect of 3’,5’-cAMP on the actin organization in plant cells we used an *Arabidopsis* actin cytoskeleton marker line that expresses a chimeric gene fusion of the second actin-binding domain of Arabidopsis Fimbrin 1 and Green Fluorescent Protein (35S:GFP-FABD) (Ketelaar et al., 2004). We focused on the hypocotyl cells of light-grown seedlings (**Figure 4a**). After a 1 h treatment with either mock or 10 μM Br-3’,5’-cAMP, we observed that the organization of actin filaments was similar; however, the filaments appeared thicker in plants treated with 10 μM Br-3’,5’-cAMP than in the mock control (**Figure 4b**). This suggests that 3’, 5’-cAMP may affects actin bundling. To further examine the effect of 3’, 5’-cAMP on actin organization we used latrunculin B (Lat B), a toxin known to bind to actin monomers in a 1:1 ratio, to prevent actin polymerization (Coue et al., 1987). A 1 h treatment with 10 μM Lat B treatment resulted in severe disruption of the actin cytoskeleton and the formation of fewer and shorter filaments. By comparison, in the combined treatment with 10 μM 3’,5’-cAMP and Lat B we observed that Br-3’,5’-cAMP mitigated the Lat B effects (**Figure 4b**). In line with the microscopic data, WT plants growing on MS medium supplemented with 1 μM Lat B showed significantly reduced hypocotyl length (**Supplementary Figure 6, Supplementary Table 10**), but this length was partially restored in plants growing simultaneously in MS medium containing 1 μM Lat B + 10 μM Br-3’,5’-cAMP (**Figure 4c,d, Supplementary Table 10**). The addition of 10 μM Br-3’,5’-cAMP alone did not affect the hypocotyl lengths of wildtype (WT) plants (**Supplementary Figure 6**).

**Figure 4.**
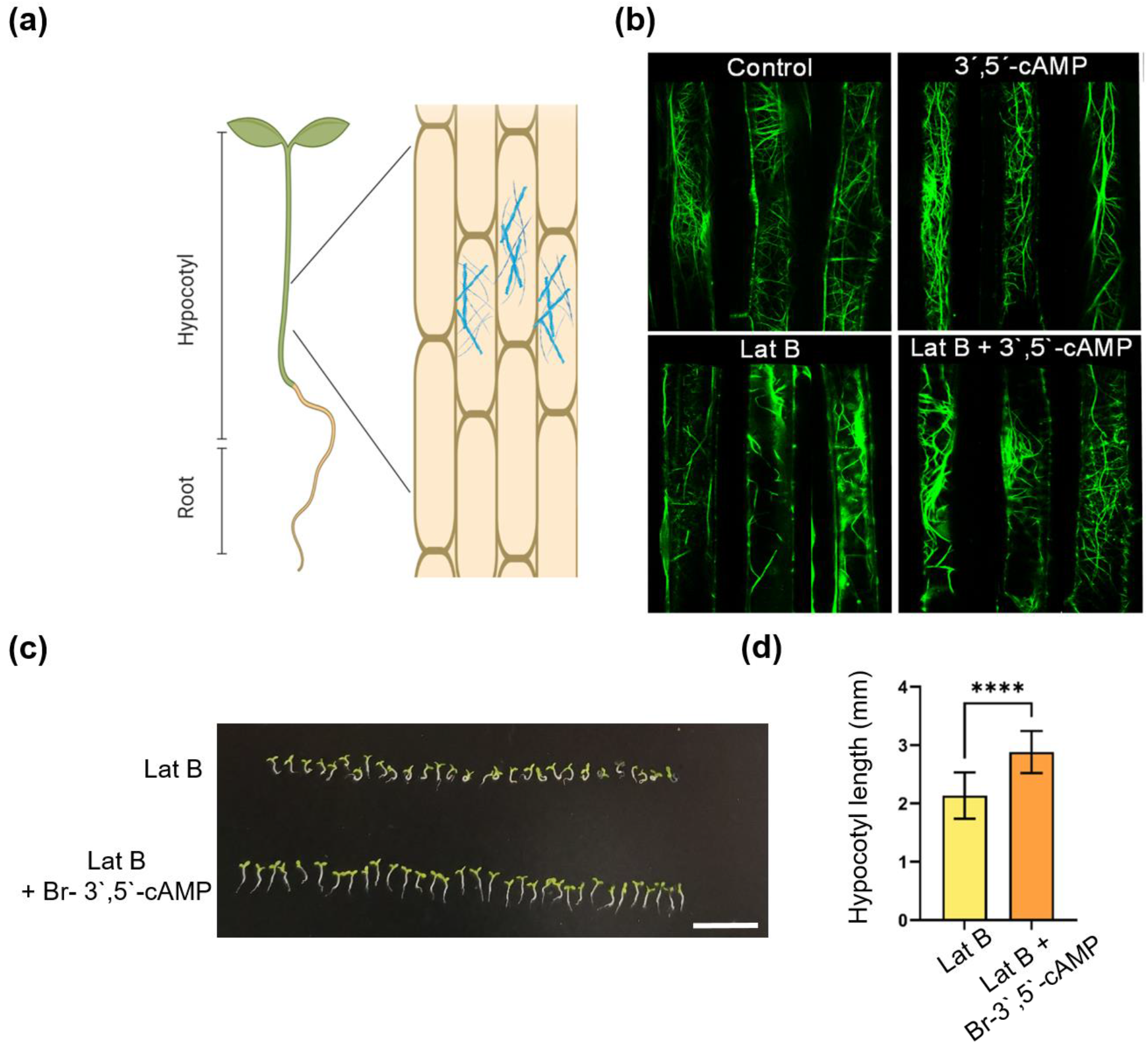
3’,5’-cAMP affects actin cytoskeleton *in planta*. (**a**) Schematic representation of the hypocotyl region that was visualized under a confocal microscope. The figure was created using BioRender.com. (**b**) Visualization of actin filaments in hypocotyl cells after 1 h treatments: control, 10 μM Br-3’,5’-cAMP, 10 μM latrunculin B (Lat B) and 10 μM Lat-B + 10 μM Br-3’,5’-cAMP treatment (room temperature). Micrographs represent three cells, each from an independent plant. (**c**) Dark grown wildtype (WT) plants growing on medium supplemented with 1 μM Lat B, or 1 μM Lat B + 10 μM Br-3’,5’-cAMP. n=29-33, Scale bar = 10 mm. WT plants growing in control conditions and on Br-3’,5’-cAMP are included in **Supplementary Figure 6**. (**d**) Hypocotyl length measurements for WT plants grown on 1 μM Lat B and 1 μM Lat B + 10 μM Br-3’,5’-cAMP. n=29-33. Student’s t-test; **** indicates p-value ≤0.0001. Measurements can be found in **Supplementary Table 10**.

### Interaction between 3’,5’-cAMP and Actin depends on other factors

To understand if the interaction between 3’,5’-cAMP and actin retrieved from TPP experiment, and later confirmed by functional analysis, is direct, we decided to use purified actin to examine actin polymerization in the presence of 3’,5’-cAMP. To this end, we used previously published protocol (Topf et al., 2017) where G-actin polymerization was measured with Microscale thermophoresis (MST), in the absence or presence of different concentrations of 3’,5’-cAMP, as well as in the presence of different concentrations of 2’,3’-cAMP (**Supplementary Table 11**). 3’,5’-cAMP did not affect the actin polymerization rate (**Figure 5a, Supplementary Figure 7**). Analogous results were obtained using pyrene actin polymerization assay and by testing thermal unfolding of G- and F-actin in the absence and presence of 3’,5’-cAMP (**Figure 5a**). Obtained results argue against a direct binding between 3’,5’-cAMP and actin. However, it has to be noted that for our assay we used mammalian actin, which in stark contrast to plant actin, can be easily purified.

**Figure 5.**
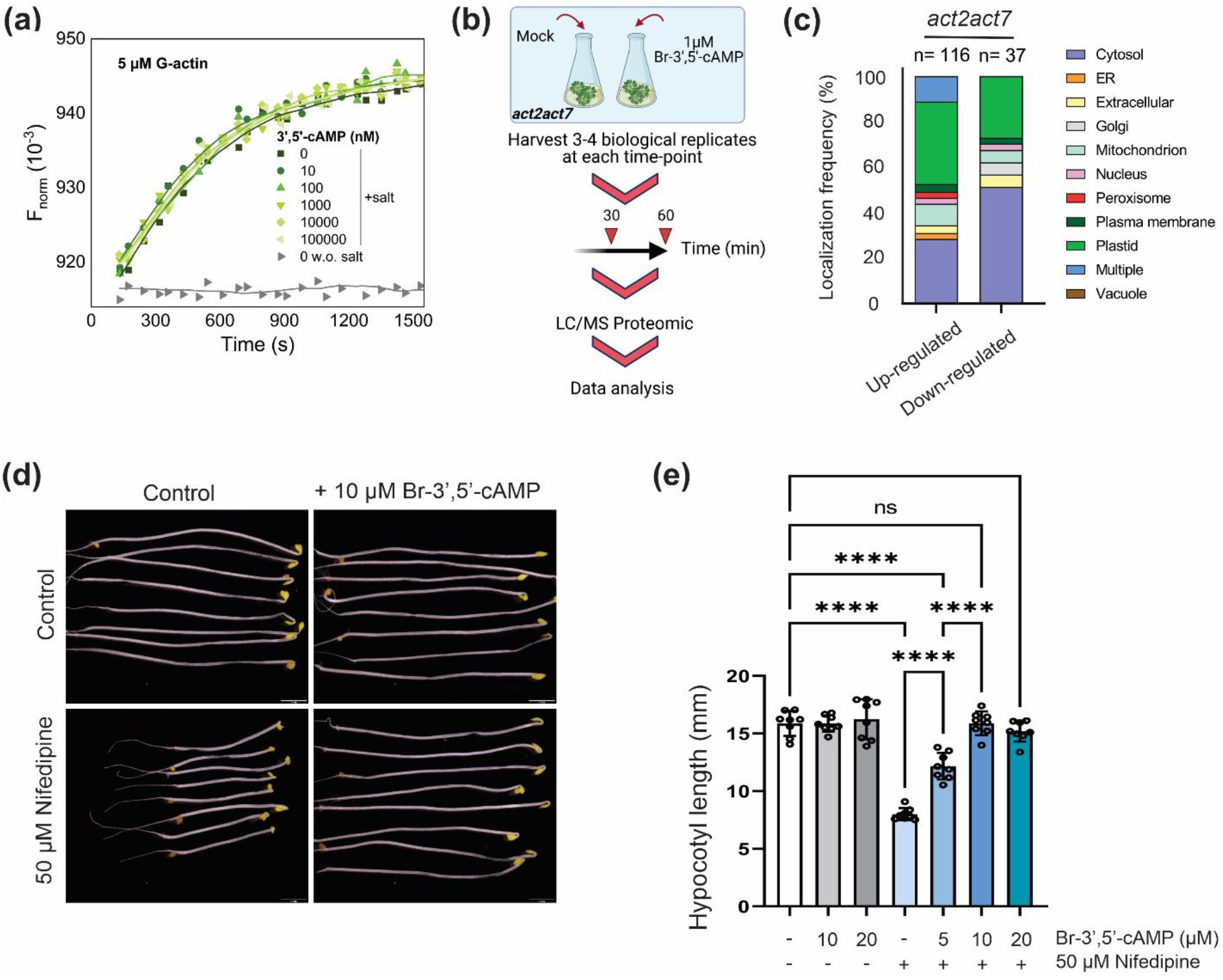
Interaction between actin and 3’,5’-cAMP depends on other factors such Ca^2+^. (**a**) MST-based actin polymerization assays with G-actin in the presence of 3’,5’-cAMP. Data sets are the sum of three experiments. (**b**) Graphical representation of the experimental design. *act2act7* seedlings grown for 10 days in liquid medium were treated with DMSO (mock) or 1μM Br-3’,5’-cAMP. Samples were collected at two different time points – 30 min and 1 h. Proteins were extracted and analyzed by LC/MS. (**c**) Localization frequency of proteins showing significant (ANOVA, p-value based on F-distribution ≤ 0.05) changes in *act2act7* seedlings in response to 0.5 and 1 h of Br-3’,5’-cAMP treatment. Data are presented as percentages for n=116 for up-, and n=37 downregulated proteins in *act2act7* plants. Color code of different compartments is presented on the legend. The whole proteomic dataset is presented in **Supplemental Table 12**. (**d**) Dark-grown WT hypocotyls on control MS medium (up left image) or on MS medium supplemented with Br-3’,5’-cAMP (up right image), or 50 μM Nifedipine (down left) or both (down right). Scale bar = 2mm. (**e**) Hypocotyl length measurements for WT plants grown on different concentrations Br-3’,5’-cAMP or Nifedipine or both. Tukey’s HSD; **** indicates p-value ≤0.0001, ns means not significant. Measurements can be found in **Supplementary Table 13**.

**Figure 6.**
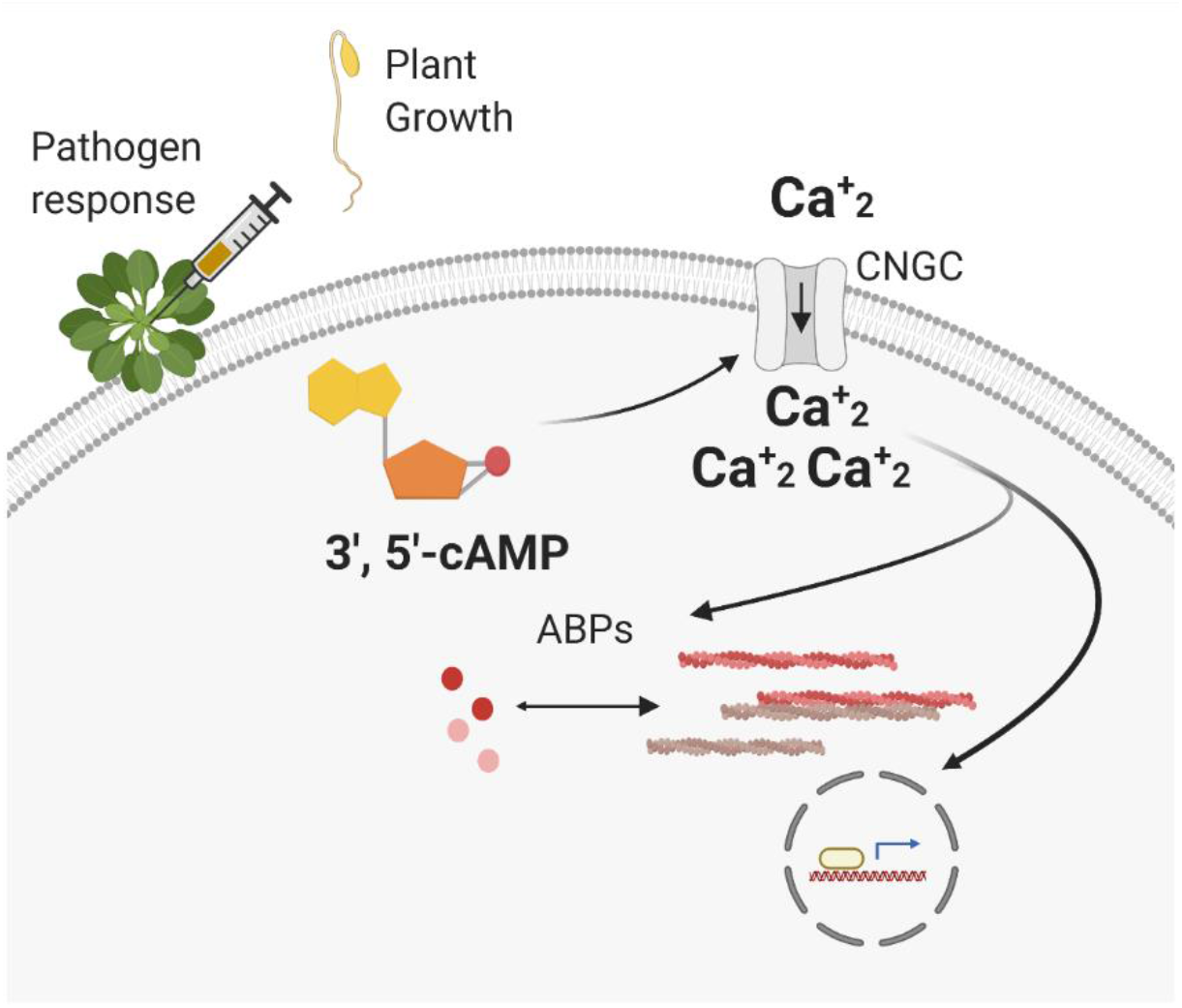
Proposed model of 3’,5’-cAMP signaling in plants. The 3’,5’-cAMP binds to cyclic nucleotide-gated cation channels (GNCGs), thereby affecting Ca^2+^ levels. Among other responses, calcium regulates gene expression as well as the actin cytoskeleton *via* the activity of the actin binding proteins (ABPs). The figure was created using BioRender.com.

### Proteomics analysis reveals a link to calcium signaling

To better understand the role of 3’,5’-cAMP in regulating actin cytoskeleton, Arabidopsis seedlings supplemented with 1 μM Br-3’,5’-cAMP, or water (control) were subjected to untargeted proteomics analysis at two time-points: 30 min and 1 h (**Figure 5b**). Rather than wild-type, we used *act2act7* mutant because of the evident phenotypic changes associated with the 3’,5’-cAMP supplementation. Proteomics analysis revealed that 3’,5’-cAMP significantly affected 153 proteins (ANOVA, *p*-value based on F-distribution ≤ 0.05), 116 being up- and 37 being down-regulated (**Supplementary Table 12**). The highest percentage of affected proteins were cytosolic and plastidial localized (**Figure 5c**). We were especially interested in proteins previously associated with the actin cytoskeleton, and notably, among the up-regulated proteins, we found ACTIN3 (AT3G53750). Although significant, the measured change was only 1.12-fold, and hence it seems unlikely the sole reason behind 3’,5’-cAMP complementation of the *act2act7* phenotype. Other proteins that stood out in the list of significantly affected proteins, are chaperonin containing TCP-1 subunit 8 (CCT8, AT3G03960), mitogen-activated protein kinase 4 (MAPK4, AT4G01370), annexin 2 (ANNAT2; AT5G65020), and plasma-membrane associated cation-binding protein 1 (PCAP1; AT4G20260) (also known as microtubule-destabilizing protein 25, MDP25) (Li et al., 2011). CCT8 is a subunit of the type II chaperonin complex, which facilitates protein folding and its known to have actin and tubulin among its well-characterized substrates (in yeast and mammalian cells) (Horwich et al., 2007; Xu et al., 2011). MPK4 has a clear role in cytoskeleton organization, since mpk24 mutant displays defective microtubule organization and is involved in phosphorylation of various microtubule-associated proteins (MAPs) (Beck et al., 2010; Mao et al., 2005; Rayapuram et al., 2018).

In plants, there is evidence that annexins may act as unconventional Ca^2+^-permeable channels, at the same time that are involved in different developmental processes, but, more interestingly, some annexins can bind F-actin and probably serving as scaffolding proteins for Ca^2+^ and the actin cytoskeleton (Qian and Xiang, 2019). On the other hand, PCAP1/MDP25 is a negative regulator of hypocotyl cell elongation by destabilizing cortical microtubules (Li *et al*., 2011) and of pollen tube elongation by severing actin filaments (Qin et al., 2014), both in a calcium-dependent manner.

The upregulation of ANNAT2 and PCAP1/MDP25 might explain the partial recovery of hypocotyl length observed in *act2act7* mutants treated with Br-3’,5’-cAMP and WT seedlings grown in the presence of Lat B, since both proteins are previously described as calcium-dependent actin-binding proteins (ABPs) (Qin *et al*., 2014). Therefore, we hypothesized, that Br-3’,5’-cAMP could be indirectly involved in the actin cytoskeleton remodeling by the upstream regulation of calcium homeostasis and the subsequent activation of ABPs.

Indeed, it has been reported that the reversible binding of cGMP, cAMP, or calmodulin (CaM) regulates CNGCs opening, and therefore cytosolic Ca^2+^ levels (Duszyn *et al*., 2019). If that is true, 3’,5’-cAMP will counteract the action of the L-type Ca^2+^ channel blocker nifedipine (De Vriese et al., 2018; Reiss and Herth, 1985). To test this, we measured hypocotyl length of Arabidopsis seedlings grown in MS medium supplemented with nifedipine and nifedipine with increasing concentrations of Br-3’,5’-cAMP. Notably, the reduction of hypocotyl length observed in seedlings treated with nifedipine was recovered in a dose-dependent manner by Br-3’,5’-cAMP (**Figure 5d,e, Supplementary Table 13**). It is remarkable that the addition of 10 μM Br-3’,5’-cAMP was enough to restore the hypocotyl length of seedling growing on medium supplemented with nifedipine; however, the addition of a higher Br-3’,5’-cAMP concentration did not produce any extra effect once the hypocotyl length was fully restored compared with the control treatment. Besides, as previously observed, 3’,5’-cAMP did not affect WT plants grown under control conditions.

## Discussion

Cyclic nucleotide-gated channels (CNGCs) are the only known direct protein targets of 3’,5’-cAMP signalling in plants (Jarratt-Barnham *et al*., 2021) known to conduct Na^+^, K^+^, and Ca^2+^ ions. To date, the reported physiological effects of 3’,5’-cAMP signaling in plants have been linked to the regulation of CNGCs (Duszyn *et al*., 2019), which transduce alterations in the intracellular concentrations of cyclic nucleotides into changes in ion concentrations.

Here, we identified ACTIN2 among putative 3’,5’-cAMP interactors using TPP. 3’,5’-cAMP supplementation increased ACTIN2 thermal stability which indicates a change in the actin interaction status. Proteins retrieved in a TPP experiment can be both direct and indirect targets of the tested ligand (Savitski *et al*., 2014). Results of the polymerization assay using purified actin argue against the direct binding. However, it has to be noted that we did use mammalian actin, which, however highly similar, is not identical to the plant actins (Kandasamy *et al*., 2012).

To challenge the functional importance of the retrieved interaction, and understand if interaction is direct or indirect, we used *act2act7* mutant. Remarkably, an increase of the 3’,5’-cAMP levels was sufficient to complement the *act2act7* short hypocotyl phenotype arguing for 3’,5’-cAMP role in the regulation of actin cytoskeleton. Along similar lines 3’,5’-cAMP supplementation was also sufficient to counter the activity of an actin-depolymerizing drug latrunculin B. A previous study showed that the disruption of actin filaments after applying Lat B or jasplakinolide resulted in mitochondrial calcium release into the cytoplasm (Wang et al., 2010). Besides, (Wang et al., 2004) reported the increase of cytoplasmic Ca2+ in Arabidopsis pollen tubes and pollen protoplasts due to the addition of actin polymerization blockers cytochalasin B and D. Remarkably, this increase is abolished by the addition of Ca2+ channel blockers. Thus, it seems that counteraction of Lat B action by application of 3’,5’-cAMP might be regulated by calcium signaling.

To unravel the mechanism by which 3’,5’-cAMP affects the actin cytoskeleton we decided to perform un-targeted proteomic experiment and see which are the proteins affected by 3’,5’-cAMP. Proteomics analysis of the *act2act7* seedling supplemented with 3’,5’-cAMP points to two different, however not exclusive, mechanisms. Modest, however, significant upregulation of ACTIN3, one of the reproductive actin isoforms, in 3’,5’-cAMP treated *act2act7* seedlings suggests that 3’,5’-cAMP might directly regulate level of actin protein. This, however, cannot explain the change in actin interaction status measured by TPP. In addition to ACTIN3, among the proteins displaying significant changes in 3’,5’-cAMP-treated *act2act7* seedlings, we found two calcium-dependent ABPs. This observation was intriguing as Ca^2+^ signaling is known to be involved in the remodeling of the actin cytoskeleton *via* binding and regulation of the actin-binding proteins and or calcium-stimulated protein kinases (Qian and Xiang, 2019).

Considering the known role of 3’,5’-cAMP in regulating Ca^2+^ flux via activation of specific members of the CNGCs family (Jarratt-Barnham *et al*., 2021) we hypothesize that 3’,5’-cAMP modulates Ca^2+^ levels, upstream of calcium-dependent ABPs. Rescue of the hypocotyl growth in the nifedepine-treated seedlings attests to the functional relationship between 3’,5’-cAMP and Ca^2+^ signaling in the regulation of cell expansion, and hypocotyl growth.

In the past 3’,5’-cAMP signalling upstream of the CNGCs and Ca^2+^ was associated with activating plant innate immune response (Ma and Berkowitz, 2011). Arabidopsis cAS lines, which have a lower level of active 3’,5’-cAMP are susceptible to the bacterial pathogen *Pseudomonas syringae* DC30000, a phenotype the authors attributed to a delay in the cytosolic calcium elevation, upstream of the redox-mediated defense mechanisms (Sabetta *et al*., 2019). The increase in actin polymerization and actin filament abundance in the epidermal cells is also a known defense mechanism against *P. syringae* (Henty-Ridilla et al., 2013). Treatment with Lat B results in susceptibility to bacterial infection (Porter and Day, 2016). Hence, and in the future, it will be interesting to investigate whether 3’,5’-cAMP signaling regulates actin cytoskeleton in response to bacterial infection.

In the dynamic network of actin filaments, microtubules, and accessory proteins that constitute the plant cytoskeleton, the necessary balance of actin filament polymerization and turnover is controlled by the competitive binding of capping proteins, destabilizers and assembly catalyzers (Carlier and Shekhar, 2017; Huang et al., 2003). In this line, the direct or indirect
 upregulation by 3’,5’-cAMP of proteins involved in growth and severing of actin filaments (and also of microtubules) may contribute to maintain the pool of polymerization-competent actin monomers and help to partially restore cytoskeletal dynamics in *act2act7* mutant, explaining the partial phenotype rescue.

A role of 3’,5’-cAMP signaling in the dynamics of the actin cytoskeleton is well documented in animal cells (Gerits *et al*., 2007) for instance in the context of an actin-dependent cell migration (Howe, 2004) and neuronal growth (Chen et al., 2003). The role of 3’,5’-cAMP signaling for regulating the actin cytoskeleton in animal cells was associated with PKA activity. PKA can directly phosphorylate monomeric actin, thereby decreasing the actin polymerization rate (Howe, 2004). Moreover, PKA can associate with and regulate the various components of the actin cytoskeleton *via* actin cytoskeleton A-kinase anchoring proteins (AKAPs) (Howe, 2004). Here we prove, that in plants, cytoskeleton regulation by 3’,5’-cAMP tightly but not exclusively depends on calcium signaling.

## Materials and methods

### Cyclic nucleotides and other compounds

The 3’,5’-cAMP (A6885), 2’,3’-cAMP (A9376), Br-3’,5’-cAMP (B5386) and Br-adenosine (B6272) were purchased from Sigma (Sigma-Aldrich, St. Louis, MO, USA). Br-2’,3’-cAMP was purchased from Biolog (Biolog Life Science Institute, Bremen, Germany). Forskolin (F6886), and theophylline (T1633) were purchased from Merck (Merck, Darmstadt, Germany). Latrunculin B (L5288) and nifedipine (N7634) were purchased from Sigma-Aldrich.

### Plant material and hypocotyl measurement

*Arabidopsis thaliana* wild-type (Wassilewskija) and double null *act2act7* mutant seeds were provided by Dr. Arun Sampathkumar. The cAS lines were kindly provided by Prof. Maria C. de Pinto and Dr. Emanuela Blanco. For time course experiments focused on hypocotyl growth, after two days of stratification, seedlings were grown on 0.5 MS agar horizontal plates supplied with 1% of sucrose without (control) or with 10 μM Br-3’,5’-cAMP for 5 days in the dark. For the experiment with forskolin, theophylline, nifedipine and Br-2’,3’-cAMP, the hypocotyls were measured 7 days after stratification. Treatments were performed as stated in the results section. The cAS lines were measured 5 days after stratification. Hypocotyls were measured using the ROI line function in the Fiji ImageJ program from images taken under a binocular light microscope. For each experiment, the number of replicates (n) (plants) is included in the figure caption.

### Thermal proteome profiling and proteomic analysis

A 3 g sample of cells from an *Arabidopsis* cell culture growing in the light were ground up with a mortar and pestle in 3 mL of lysis buffer (Tris HCl pH 7.5 50 mM, NaCl 500 mM, MgCl_2_ 1.5 mM, DTT 5 mM, NaF 1 mM, PIC 100×, Na_3_VO_4_ 0.1 mM, PMSF 1 mM). The lysates were centrifuged at 35000 rpm for 1 h at 4°C, followed by a 20 min centrifugation at 4°C and 4000 rpm using 10-kD Amicon^®^Ultra centrifuge filter (Merck, Darmstadt, Germany) to remove free (proteinun bound) small molecules, such 3’,5’-cAMP. The resulting clear soluble protein extract was incubated with 10 μM 3’,5’-cAMP or a DMSO control for 30 min at room temperature. For further steps, the protocol by Franken *et al* (Franken *et al*., 2015) was followed with adaptation to a full protein extract from an Arabidopsis cell culture. In brief, a 3 min temperature treatment was performed using an Eppendorf PCR machine and a gradient of different temperatures was applied: RT, 39,9, 45,8, 51,1, 56,2, 60,1, 65,6, 70,9, 76,0, and 79,9 °C. The samples were then centrifuged for 1 min at RT at 14000 rpm to remove precipitated proteins from the solution. The supernatant was collected into a new tube and the proteins were precipitated with ice-cold acetone (1/4) and dried for 2–4 h in a centrifugal evaporator. The proteins were then prepared for mass spectrometry analysis.

### Protein preparation

Dried protein pellets were resuspended in 30 μL of urea buffer (6 M urea and 2 M thiourea in 40 mM ammonium bicarbonate). The amount of protein for trypsin digestion was quantified based on a control – RT sample where protein content is the highest. Approximately 3 μL (corresponding to 50 μg of total protein content in the RT sample) was used for protein digestion. Cysteine residues were reduced by the addition of 2.5 μL 100 μM DTT for 30 min at RT, followed by alkylation with 2.5 μL 300 mM iodoacetamide for 20 min in the dark. The proteins were enzymatically digested using LysC/Trypsin Mix (Promega) according to the manufacturer’s instructions. The peptides were then desalted on C18 SepPack columns, dried to approximately 4 μL using a centrifugal evaporator, and stored at –80°C until measurement.

### Mass spectrometry analysis

Desalted peptides were suspended in 146 μL of the loading buffer (2% ACN, 0.2% TFA) and 3 μL (corresponding to 1 μg of peptides in RT sample) were loaded onto a C18 reversed-phase column connected to an Acquity UPLC M-Class system. The sample was fractionated using a 155 min gradient at a flow rate of 400 μL min^-1^. Proteins were eluted with a gradient of acetonitrile in 0.1% acetic acid as follows: 0 to 14 min: 3.2% ACN; 14 to 34 min: 3.2–7.2% ACN; 34 to 104 min: 7.2–24.8% ACN; 104 to 134 min: 24.8–35.2% ACN; 134 to 135 min: 35.2–76% ACN; 135 to 140 min: 76% ACN; 140 to 141 min: 76–3.2% ACN; 141 to 155 min: 3.2% ACN. Mass spectra were acquired using a Thermo Q Exactive HF operated with a data-dependent method with following settings: resolution set to 60,000, scan range from 300.0 to 1600.0 m/z, maximum fill time of 50 ms, and an AGC target value of 3e6 ions. A maximum of 10 data-dependent MS2 scans were performed in the ion trap set to an AGC target of 1e5 ions, with a maximal injection time of 80 ms. Precursor ion fragmentation was achieved with collision-induced fragmentation with a normalized collision energy of 27 and isolation width of 1.6 m/z. Charge states of 1 and ≥8 were rejected. The obtained chromatograms were processed using MaxQuant version 1.5.2.8 (Jürgen Cox & Mann, 2008). Peptides were identified using the built-in search engine, Andromeda (Jurgen Cox *et al*, 2011) with an *Arabidopsis thaliana* proteome library, modified in December 2016 and downloaded from Uniprot (http://www.uniprot.org/proteomes/UP000002311). A contaminant database was also included in the search. Detailed MaxQuant settings and parameters are shown in **Supplementary Table 1**. A total of 5245 protein groups were initially identified in samples treated with the temperature gradient. The number of identified protein groups decreased to 4800 when contaminants, decoy hits, and proteins quantified with less than two unique peptides were filtered out.

### Thermal proteome profiling (TPP) analysis

Proteins with significant differences between both experimental conditions were detected by nonparametric analysis of response curves (Childs *et al*., 2019). Only proteins quantified with at least two unique peptides in all three replicates were considered. If a protein exhibited zero intensities at all temperatures in an experimental group, it was omitted from the analysis. In total, 4862 proteins passed these filters and were included in the analysis. The sample representing the control group of replicate 1 at 70.9 °C was excluded from further analysis because exploratory data analysis showed a systematically altered distribution shape of measurements in this sample. Raw intensity values were converted to log2 scale and normalized by centering them to a common median per temperature. For each protein, the values were then scaled to the condition-specific mean of the corresponding values at the lowest temperature. Finally, the scaled and normalized values were transformed to a non-negative range by applying the function *f*(*x*) = *log_2_*(2^*x*^ + 1) to each value.

The effects of 3’,5’-cAMP treatment were assessed by fitting two separate models per protein (Childs *et al*., 2019). The null model assumed that the melting curves in both conditions were explained by the same sigmoid mean function

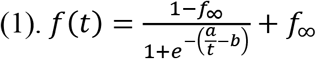

with temperature t, shape parameters a, b, and a lower horizontal asymptote *f*_∞_. The alternative model assumed that they were explained by condition-specific functions according to Eq. (1).

Improvements in goodness-of-fit between null and alternative model were then quantified for each protein with an F-statistic

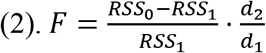

Here, RRS_0_ and RRS_1_ are the sums of squared residuals of the null and alternative models for a particular protein, and d_2_ and d_1_ are the distribution parameters of the *F*-distribution, estimated as described in (Childs *et al*., 2019).

For each protein, a p-value was computed by comparing its *F*-statistic to an *F*-distribution with the estimated parameters. All p-values were adjusted for multiple testing by Benjamini-Hochberg correction (Benjamini and Hochberg, 1995). *T_m_* values of the alternative model per condition were calculated as described in (Franken *et al*., 2015), and *T*_m_ - differences between both treatment groups were computed from these values.

### Western blotting for actin stability

Western blots were prepared from lysates from *Arabidopsis* rosettes incubated with or without 3’,5’-cAMP (as in the TPP experiment). Each lysate was divided into four parts for different temperature treatments (50, 52, 55, 60 °C). The protein samples were separated by SDS-PAGE and blotted onto membranes. Immunoblot detection was performed with specific antibodies using the enhanced chemiluminescence system (GE Healthcare). Signals were detected with the G:box Chemi Imaging System (Syngene). Antibodies used for the western blots were obtained from commercial suppliers: anti-actin monoclonal antibody (Sigma-Aldrich A0480-200μL) and anti-mouse antibody (Agrisera AS111772). Western blot quantification was performed with the open-source Fiji software.

### Confocal microscopy - actin visualization and analysis

Actin filaments were observed with a Leica DM6000B/SP5 confocal laser-scanning microscope (Leica Microsystems, Wetzlar, Germany) in *Arabidopsis* seedlings expressing the FABD protein-domain known to bind to actin (P35S:GFP-FABD). The GFP-FABD signal was visualized using 488 nm laser excitation and emission between 500 and 520 nm.

### Microscale thermophoresis (MST), pyrene-actin assay, and thermal protein unfolding

Actin was prepared from chicken skeletal muscle as described (Franz et al., 2020). MST-based actin polymerization was carried out using the Monolith NT.115 (Nanotemper, Munich, Germany) as described (Topf *et al*., 2017) with the following modifications: instead of ATTO488, NTred-maleimid 2^nd^ generation was used for actin labeling at 3-fold molar excess. The fraction of fluorescently labeled G-actin vs total G-actin concentration was 5%. IR-laser power was set to 40% and the red LED power was adjusted between 4% and 12% dependent on the experiment. The pyrene-actin polymerization assay was performed as described (Walter et al., 2020) using Tris-HCl buffer (50 mM Tris-HCl, 50 mM KCl, 2 mM MgCl_2_, pH=7.4 at room temperature) instead of imidazole buffer for initiating polymerization. cAMP binding assays to G-actin were performed with the Monolith NT.115 (Nanotemper, Munich, Germany) using 50 nM NTred-maleimid 2^nd^ generation labelled G-actin and increasing concentrations of cAMP. IR-laser power was 40% and the red LED power 12%. All measurements were performed at 25 °C. Thermal unfolding of G-actin (2 μM) and F-actin (2 μM) was performed with the Tycho NT.6 (Nanotemper, Munich, Germany) according to the manufacturer’s instructions in G-buffer (5 mM Tris-HCl, 0.2 mM CaCl_2_, 0.2 mM ATP, 0.5 mM DTT, 0.02 % w/v NaN_3_, pH = 8.0 at 4 °C) or Tris-HCl polymerization buffer, in the absence and presence of nucleotides.

### Feeding experiment

One point five milligrams of *act2act7* mutant seeds were grown in 30 ml of 0.5 liquid MS supplied with 1% sucrose at 120 rpm on an orbital shaker. After 11 days of growth, the medium was exchanged for fresh medium and incubated for another 3 days. At this point, 1 μM Br-3’,5’-cAMP for 0.5 or 1 h was applied, using water as control. Samples were harvested after treatment, briefly dried on a paper towel and immediately frozen with liquid nitrogen until use.

### Protein extraction after feeding experiments

One hundred milligrams of ground frozen tissue were mixed and vortexed with 350 μl of cold lysis buffer (Tris HCl pH 7.5 50 mM, NaCl 500 mM, MgCl_2_ 1.5 mM, DTT 5 mM, NaF 1 mM, PIC 100×, Na_3_VO_4_ 0.1 mM, PMSF 1 mM). The lysate was centrifuged at 12,000 rpm for 5 min at 4 °C, then, 300 μl of supernatant were transferred to a new tube and mixed 3:1 with cold acetone. The pellet was precipitated by centrifugation at max speed for 1 min at 4 °C and dried in a centrifugal evaporator. The protein pellets were prepared for MS as described in section “Protein preparation”, MaxQuant setting for that experiment are shown in Supplementary Table 2.

### Proteomic data availability

The mass spectrometry proteomics data from TPP experiment have been deposited to the ProteomeXchange Consortium via the PRIDE (Perez-Riverol et al., 2019) partner repository with the dataset identifier PXD019537. Proteomic data from feeding experiment are in the process of submission.

## Acknowledgments

We thank Dr. Rene Schneider, Dr. Sakutaru Kijima, Prof. Taro Uyeda, Prof. Jakub Sedzinski, and Prof. Lothar Willmitzer for valuable scientific input. We are grateful to Prof. Maria C. de Pinto and Dr. Emanuela Blanco for sharing their cAMP sponge lines with us and to Dr. Michal Gorka for the input on proteomics. We are also grateful to KAUST core lab for running samples in their proteomic facility.

## Author Contributions

A.Sk. conceived the project; A.Sk., M.C., T.G., and A.Sa. designed experimental strategy; P.F., performed Microscale thermophoresis (MST), pyrene-actin assay, and thermal protein unfolding for actin protein; M.L. run proteomic samples from TPP experiment, D.C. analyzed the TPP data; J.C.M. provided technical assistance for the TPP; A.M. provided technical assistance for the hypocotyl experiments; M.S provided help and organized supplementary material, N.E.F performed feeding experiment of *act2act7* seedlings with 3’,5’-cAMP, prepared samples for proteomic, analyzed the data, and performed phenotypical experiments with nifedipine, M.C. performed the majority of the experimental work; A.Sk. and M.C. wrote the manuscript with input from N.E.F.

## Supplementary Tables

**Supplementary Table 1. MaxQuant output parameters.txt file. It contains parameters used for processing RAW chromatograms of WT+Br-3’,5’-cAMP dataset**

**Supplementary Table 2. MaxQuant output paramters.txt file. It contains parameters used for processing RAW chromotograms of *act2act7* + Br-3’,5’-cAMP dataset**.

**Supplementary Table 3. Raw proteomic data from thermal proteome profiling (TPP).**

**Supplementary Table 4. Proteomics data analyzed by thermal proteome profiling**.

**Supplementary Table 5.Western blot quantification form Figure 1d**.

**Supplementary Table 6. Hypocotyl length measurements from the Br-3’,5’-cAMP WT and act2act7 feeding experiment**.

**Supplementary Table 7. Hypocotyl length measurements of *act2act7* mutant growing on forskolin, theophylline and Br-2’,3’-cAMP**.

**Supplementary Table 8. Hypocotyl length measurements from the Br-3’,5’-cAMP (different concentrations) act2act7 feeding experiment. For Figure 2d.**

**Supplementary Table 9**.

**Hypocotyl length measurements of 3’,5’-cAMP sponge lines treated with Br-3’,5’-cAMP**.

**Supplementary Table 10. Hypocotyl length measurements from wildtype (WT) plants growing on control medium or supplemented with Br-3’,5’-cAMP, latrunculin B (Lat B), or Lat B plus Br-3’,5’-cAMP**.

**Supplementary Table 11. Data from microscale thermophoresis (MST), pyrene-actin assay and thermal protein unfolding for Figure 5a**.

**Supplementary Table 12. Proteomic data from Br-3’,5’-cAMP feeding experiment**.

**Supplementary Table 13. Hypocotyl length measurements for WT plants grown on different concentrations Br-3’,5’-cAMP, nifedipine or nifedipine + Br-3’,5’-cAMP**.

---

**Supplementary Figure 1.**
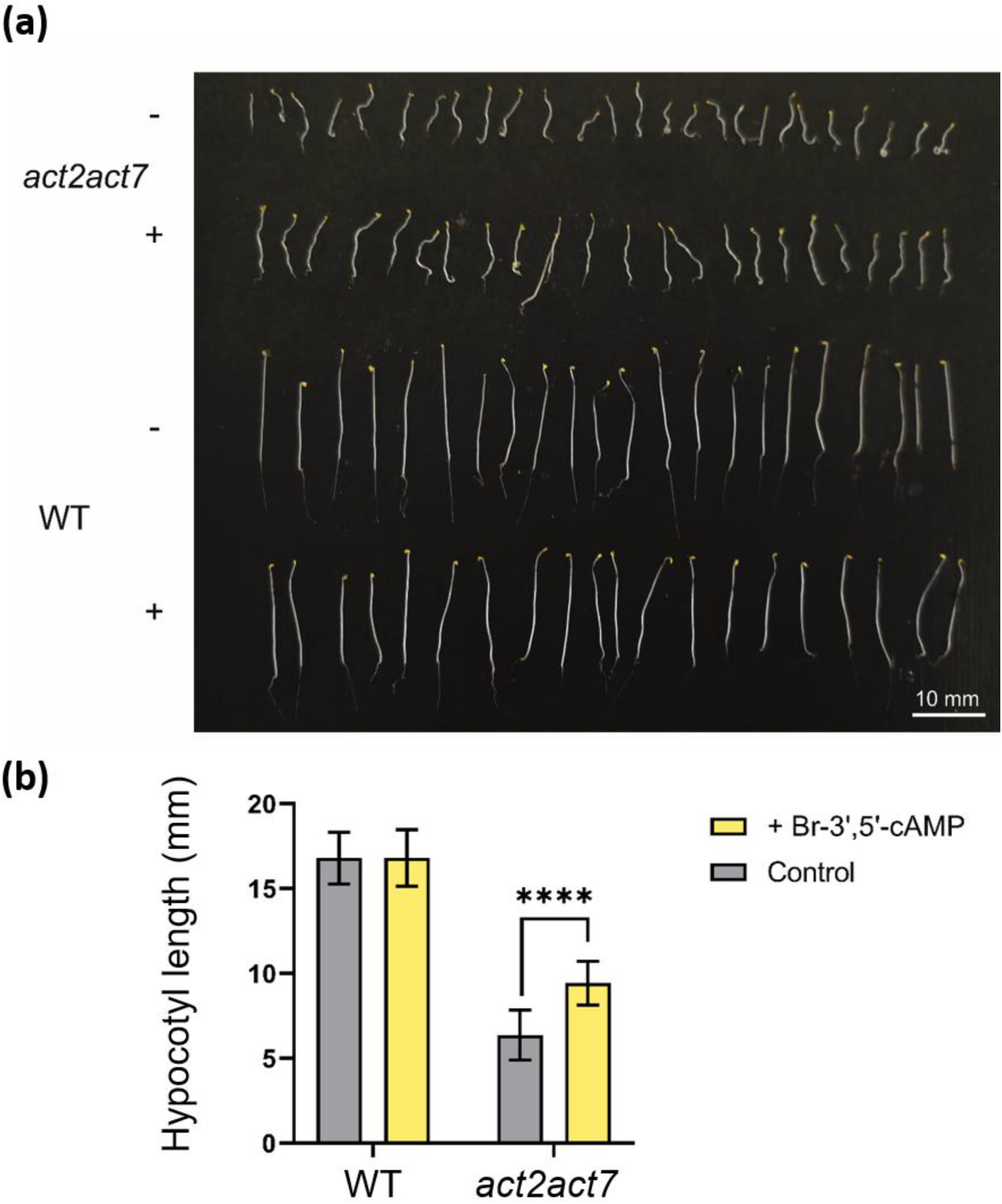
Dark grown hypocotyl length in the presence of 10 μM Br-3’,5’-cAMP. (**a**) 5-day old Arabidopsis plants growing in the dark on control MS plates without (-) or with (+) 10 μM Br-3’,5’-cAMP. Scale bar = 10 mm. (**b**) Measurement of hypocotyl growth expressed in mm. Data presented as mean ± SD, n = 22-26. Student’s T-test, **** indicates p-value ≤0.0001.

**Supplementary Figure 2.**
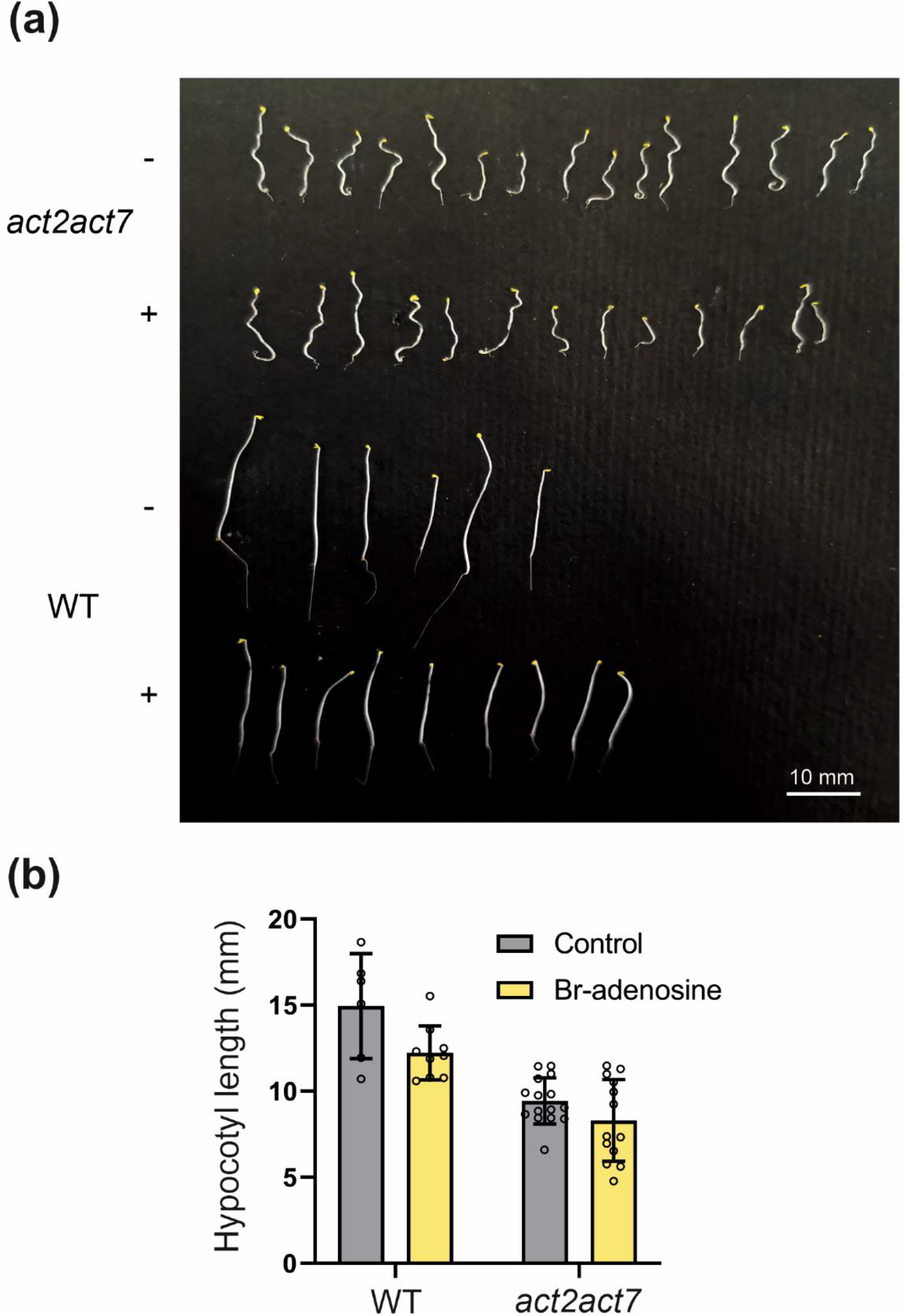
Hypocotyl growth in the presence of 10 μM Br-adenosine. (**a**) 5-day old WT and *act2act7* plants growing in the dark on control MS plates without (-) or with (+) 10 μM Br-adenosine. Scale bar = 10 mm. (**b**) Hypocotyl length of WT and *act2act7* plants grown on control or Br-adenosine supplemented medium. Hypocotyl length is expressed in mm.

**Supplementary Figure 3.**
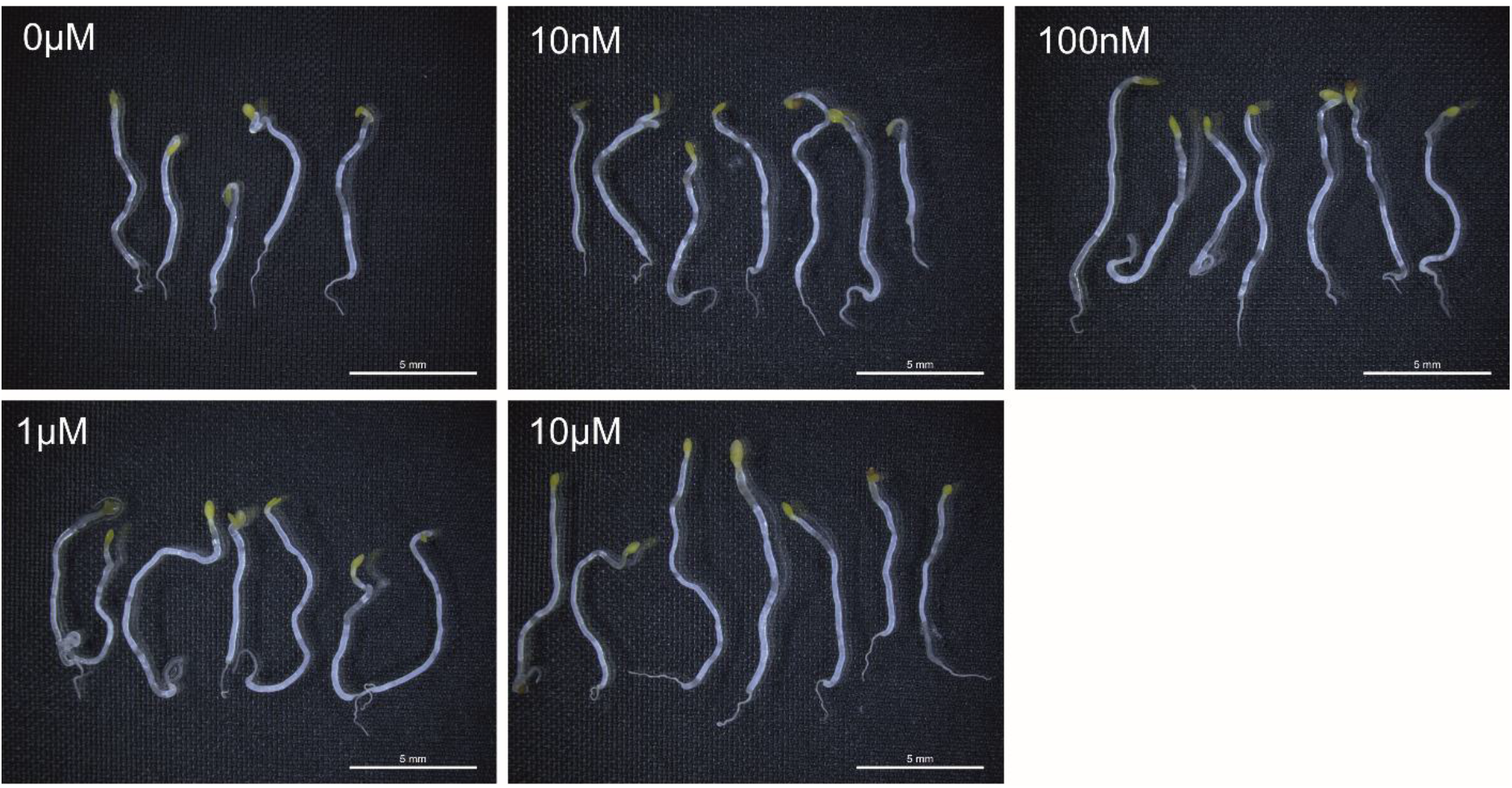
*act2act7* hypocotyl after 5 days of growth on MS medium supplemented with different concentrations of Br-3’,5’-cAMP. See also Supplementary Table S5 and Figure 2d.

**Supplementary Figure 4.**
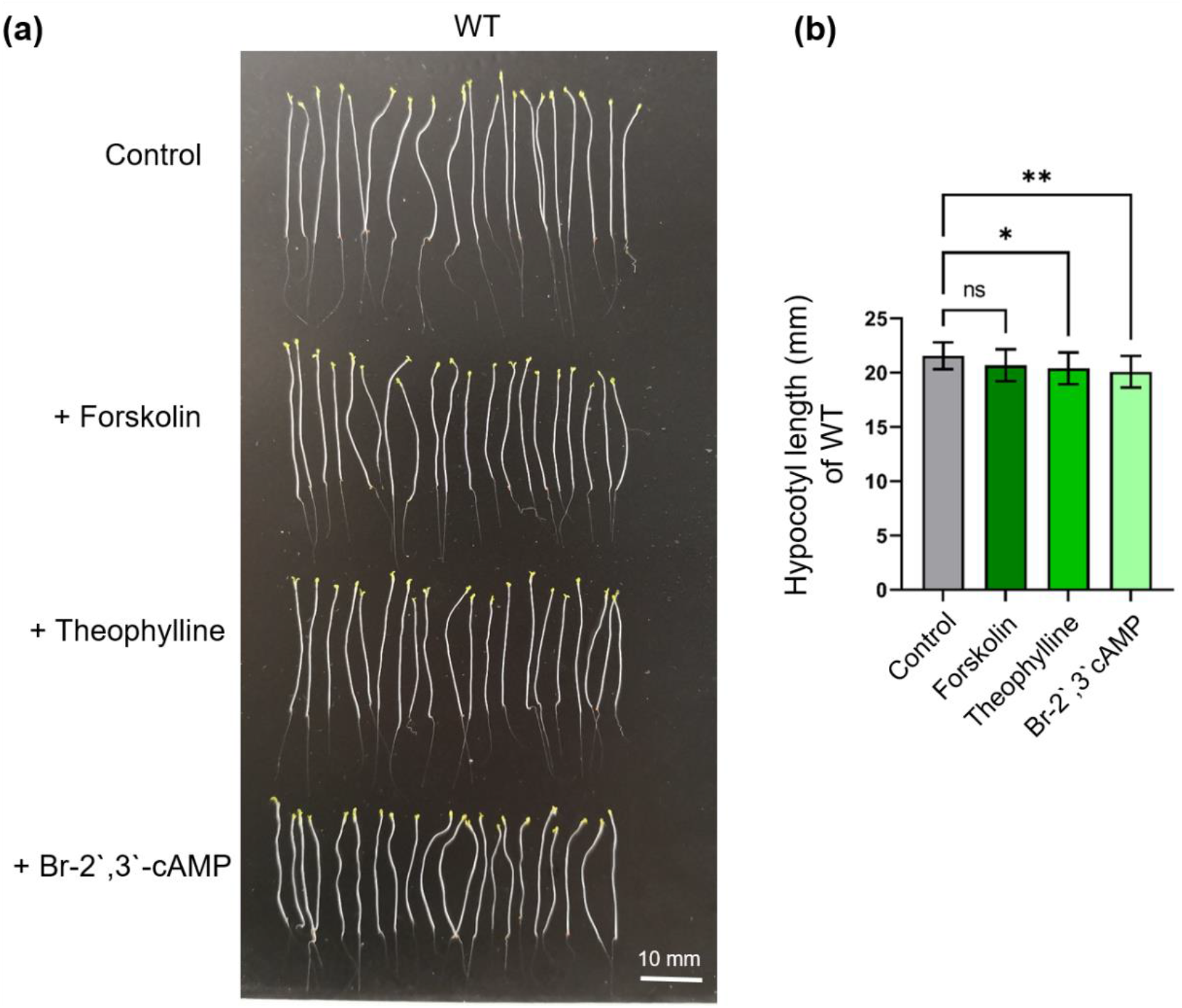
WT plants growing on MS medium without (control) or with 10 μM Forskolin, 100 μM Theophylline or 10 μM Br-2’,3’-cAMP for 5 days. (**a**) Dark grown hypocotyls. n=20, scale bar = 10 mm. (**b**) Hypocotyl length is expressed in mm. Student’s T-test, * indicates p-value ≤0.05, ** indicates p-value ≤0.01, and ns indicates p > 0.05.

**Supplementary Figure 5.**
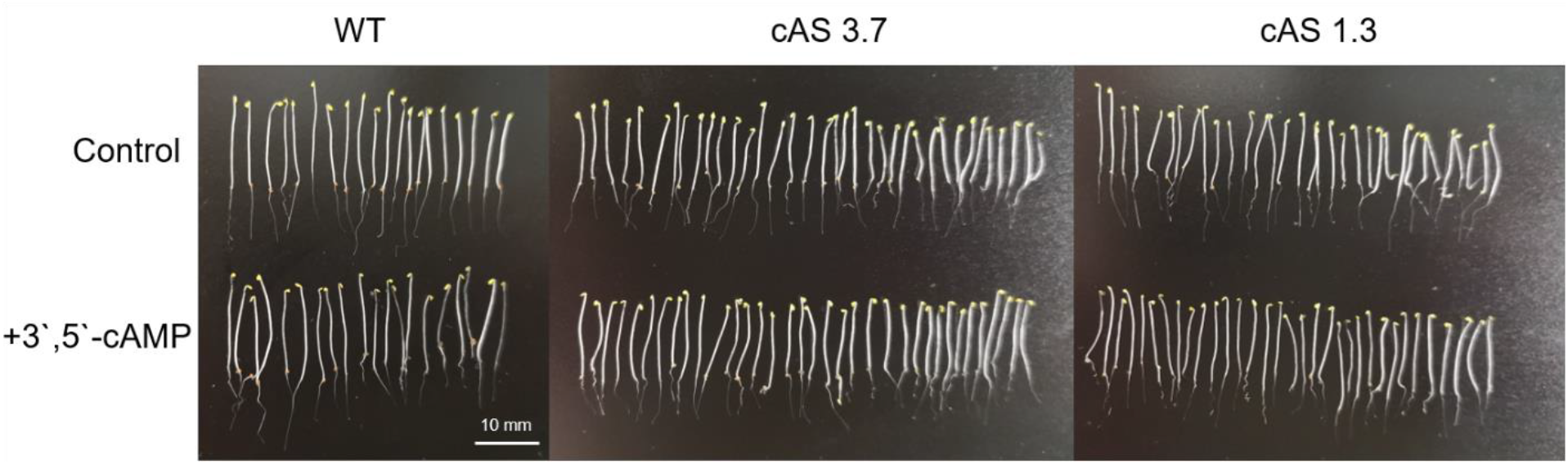
Dark grown hypocotyl of WT and cAS lines growing on control medium (Control) or supplemented with Br-3’,5’-cAMP for 5 days. n=20 for WT, n=32-36 for cAS lines. Scale bar = 10mm.

**Supplementary Figure 6.**
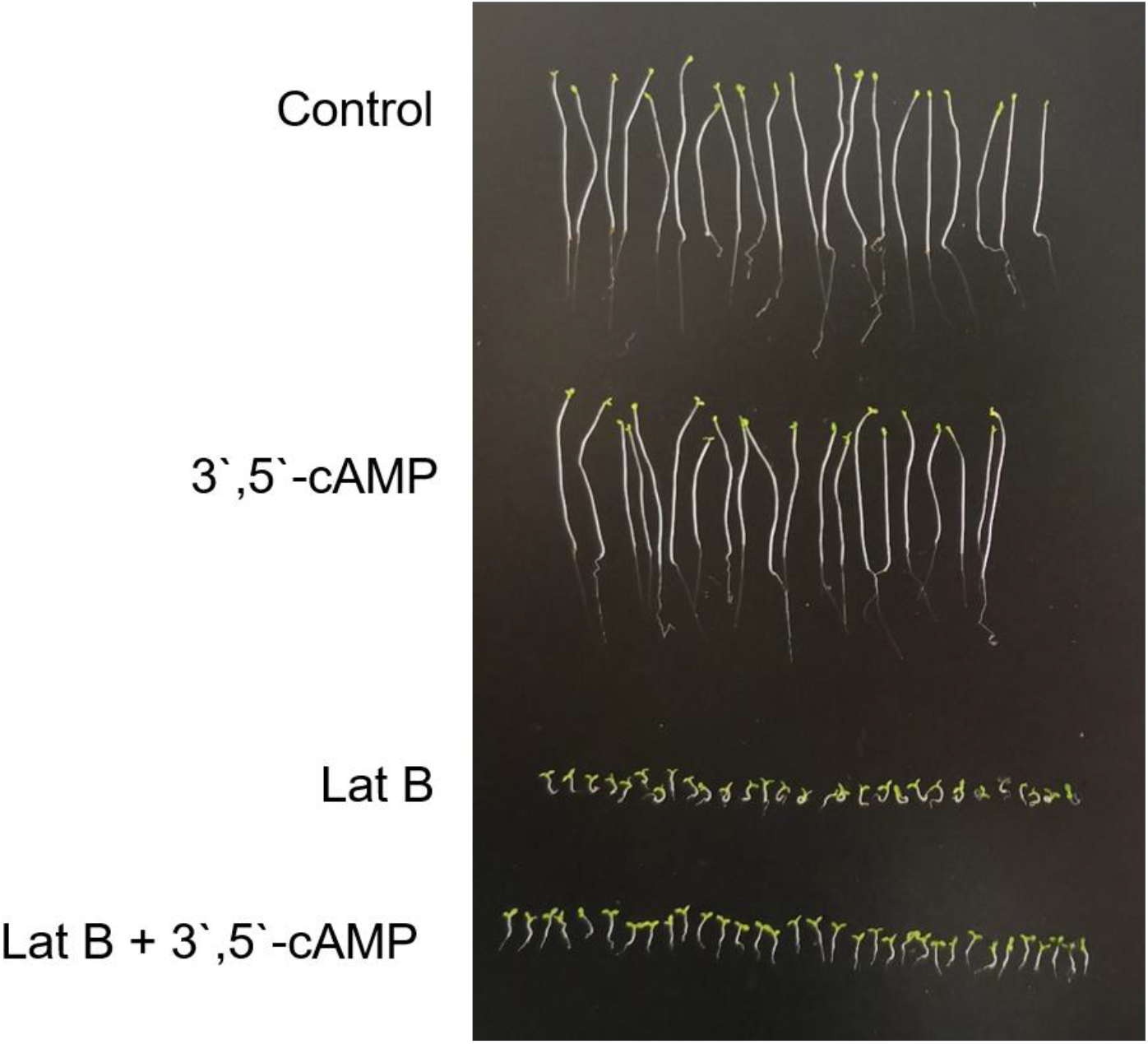
(a) Dark-grown hypocotyl length of WT Arabidopsis seedlings on MS (control) medium supplemented with 10 μM 3’,5’-cAMP, 1 μM Lat B or 1μM Lat B + 10 μM Br-3’,5’-cAMP. n= 20 for control and Br-3’,5’-cAMP, n=29-33 for Lat B and Lat B + 3’,5’-cAMP.

**Supplementary Figure 7.**
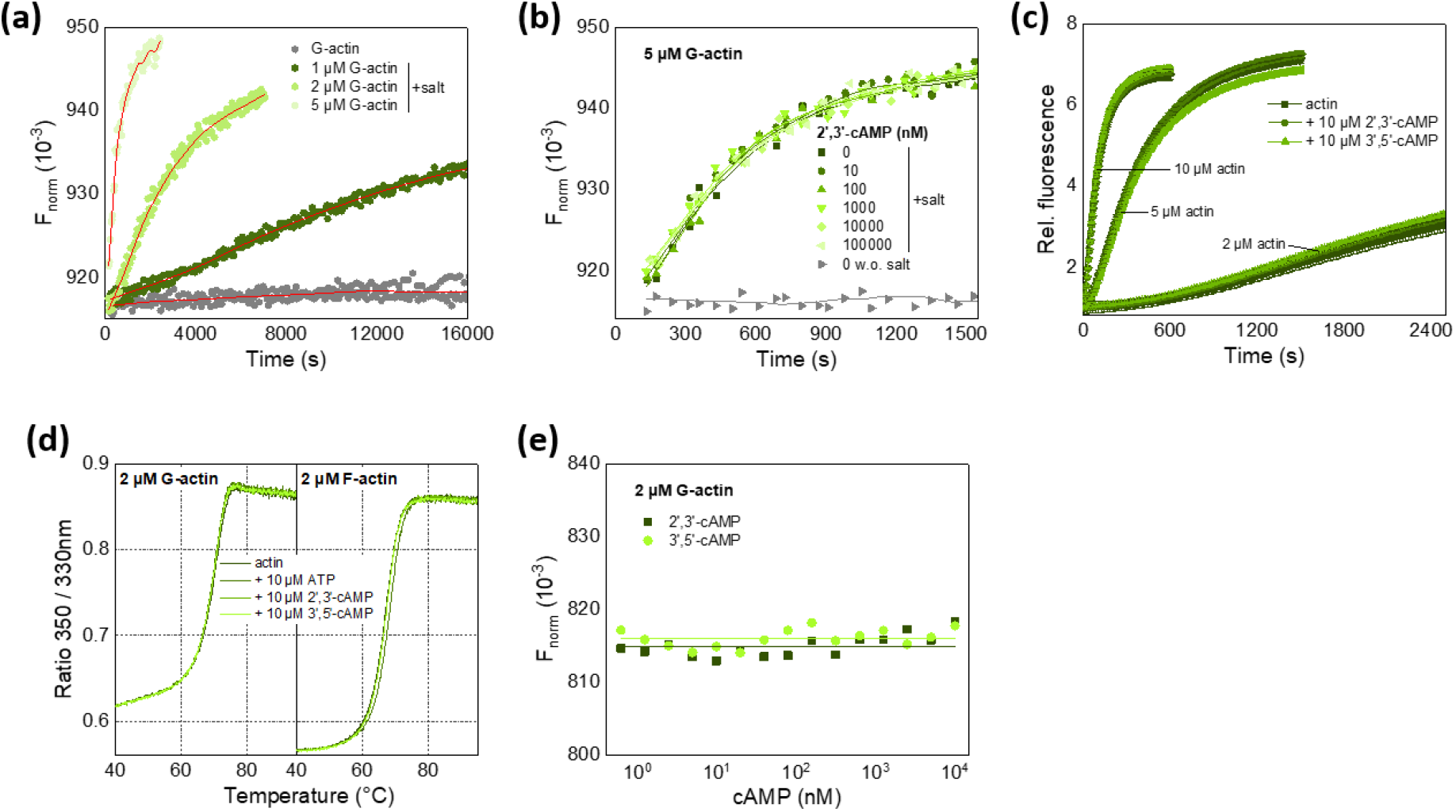
Functional assays with actin in the absence and presence of cAMP. (a) MST-based actin polymerization assays with G-actin (1, 2, 5 μM) in the absence of cAMP, and (b) in the presence of 2’,3’-cAMP. Data sets are the sum of three experiments. (c) Pyrene-actin polymerization assay with different G-actin concentrations (2, 5, 10 μM) in the presence and absence of 10 μM 2’,3’-cAMP or 3’,5’-cAMP. Data are average of three measurements ± s.d. (d) Thermal unfolding of G-actin and F-actin in the absence and presence of 10 μM 2’,3’-cAMP, 3’,5’-cAMP, or ATP. (e) MST titration experiments of G-actin with cAMP.

## Notes

### Competing Interest Statement

The authors have declared no competing interest.

## References

Alqurashi, M., Gehring, C., and Marondedze, C. (2016). Changes in the Arabidopsis thaliana Proteome Implicate cAMP in Biotic and Abiotic Stress Responses and Changes in Energy Metabolism. Int J Mol Sci 1710.3390/ijms17060852.

Amrhein, N. (1977). Current Status of Cyclic-Amp in Higher-Plants. Annual Review of Plant Physiology and Plant Molecular Biology 28:123–132. DOI 10.1146/annurev.pp.28.060177.001011.

Ashton, A.R., and Polya, G.M. (1978). Cyclic adenosine 3’:5’-monophosphate in axenic rye grass endosperm cell cultures. Plant Physiol 61:718–722. 10.1104/pp.61.5.718.

Aye, T.T., Mohammed, S., van den Toorn, H.W., van Veen, T.A., van der Heyden, M.A., Scholten, A., and Heck, A.J. (2009). Selectivity in enrichment of cAMP-dependent protein kinase regulatory subunits type I and type II and their interactors using modified cAMP affinity resins. Mol Cell Proteomics 8:1016–1028. 10.1074/mcp.M800226-MCP200.

Balague, C., Lin, B., Alcon, C., Flottes, G., Malmstrom, S., Kohler, C., Neuhaus, G., Pelletier, G., Gaymard, F., and Roby, D. (2003). HLM1, an essential signaling component in the hypersensitive response, is a member of the cyclic nucleotide-gated channel ion channel family. Plant Cell 15:365–379. 10.1105/tpc.006999.

Beck, M., Komis, G., Muller, J., Menzel, D., and Samaj, J. (2010). Arabidopsis homologs of nucleus- and phragmoplast-localized kinase 2 and 3 and mitogen-activated protein kinase 4 are essential for microtubule organization. Plant Cell 22:755–771. 10.1105/tpc.109.071746.

Bianchet, C., Wong, A., Quaglia, M., Alqurashi, M., Gehring, C., Ntoukakis, V., and Pasqualini, S. (2019). An Arabidopsis thaliana leucine-rich repeat protein harbors an adenylyl cyclase catalytic center and affects responses to pathogens. J Plant Physiol 232:12–22. 10.1016/j.jplph.2018.10.025.

Brown, E.G., and Newton, R.P. (1981). Cyclic-Amp and Higher-Plants. Phytochemistry 20:2453–2463. Doi 10.1016/0031-9422(81)83071-3.

Carlier, M.F., and Shekhar, S. (2017). Global treadmilling coordinates actin turnover and controls the size of actin networks. Nat Rev Mol Cell Biol 18:389–401. 10.1038/nrm.2016.172.

Chen, N., Furuya, S., Shinoda, Y., Yumoto, M., Ohtake, A., Sato, K., Doi, H., Hashimoto, Y., Kudo, Y., and Higashi, H. (2003). Extracellular carbohydrate-signal triggering cAMP-dependent protein kinase-dependent neuronal actin-reorganization. Neuroscience 122:985–995. 10.1016/j.neuroscience.2003.08.042.

Cheng, X., Ji, Z., Tsalkova, T., and Mei, F. (2008). Epac and PKA: a tale of two intracellular cAMP receptors. Acta Biochim Biophys Sin (Shanghai) 40:651–662. 10.1111/j.1745-7270.2008.00438.x.

Childs, D., Bach, K., Franken, H., Anders, S., Kurzawa, N., Bantscheff, M., Savitski, M.M., and Huber, W. (2019). Nonparametric Analysis of Thermal Proteome Profiles Reveals Novel Drug-binding Proteins. Mol Cell Proteomics 18:2506–2515. 10.1074/mcp.TIR119.001481.

Coue, M., Brenner, S.L., Spector, I., and Korn, E.D. (1987). Inhibition of actin polymerization by latrunculin A. FEBS Lett 213:316–318. 10.1016/0014-5793(87)81513-2.

Daly, J.W. (1984). Forskolin, adenylate cyclase, and cell physiology: an overview. Adv Cyclic Nucleotide Protein Phosphorylation Res 17:81–89.

de Rooij, J., Zwartkruis, F.J., Verheijen, M.H., Cool, R.H., Nijman, S.M., Wittinghofer, A., and Bos, J.L. (1998). Epac is a Rap1 guanine-nucleotide-exchange factor directly activated by cyclic AMP. Nature 396:474–477. 10.1038/24884.

De Vriese, K., Costa, A., Beeckman, T., and Vanneste, S. (2018). Pharmacological Strategies for Manipulating Plant Ca(2+) Signalling. Int J Mol Sci 1910.3390/ijms19051506.

Donaldson, L., Meier, S., and Gehring, C. (2016). The arabidopsis cyclic nucleotide interactome. Cell Communication and Signaling 14ARTN 10 10.1186/s12964-016-0133-2.

Dryden, W.F., Singh, Y.N., Gordon, T., and Lazarenko, G. (1988). Pharmacological elevation of cyclic AMP and transmitter release at the mouse neuromuscular junction. Can J Physiol Pharmacol 66:207–212. 10.1139/y88-036.

Duszyn, M., Swiezawska, B., Szmidt-Jaworska, A., and Jaworski, K. (2019). Cyclic nucleotide gated channels (CNGCs) in plant signalling-Current knowledge and perspectives. J Plant Physiol 241:153035. 10.1016/j.jplph.2019.153035.

Ehsan, H., Reichheld, J.P., Roef, L., Witters, E., Lardon, F., Van Bockstaele, D., Van Montagu, M., Inze, D., and Van Onckelen, H. (1998). Effect of indomethacin on cell cycle dependent cyclic AMP fluxes in tobacco BY-2 cells. FEBS Lett 422:165–169. 10.1016/s0014-5793(97)01610-4.

Franken, H., Mathieson, T., Childs, D., Sweetman, G.M., Werner, T., Togel, I., Doce, C., Gade, S., Bantscheff, M., Drewes, G., et al. (2015). Thermal proteome profiling for unbiased identification of direct and indirect drug targets using multiplexed quantitative mass spectrometry. Nat Protoc 10:1567–1593. 10.1038/nprot.2015.101.

Franz, P., Gassl, V., Topf, A., Eckelmann, L., Iorga, B., and Tsiavaliaris, G. (2020). A thermophoresis-based biosensor for real-time detection of inorganic phosphate during enzymatic reactions. Biosens Bioelectron 169:112616. 10.1016/j.bios.2020.112616.

Gao, F., Han, X., Wu, J., Zheng, S., Shang, Z., Sun, D., Zhou, R., and Li, B. (2012). A heat-activated calcium-permeable channel--Arabidopsis cyclic nucleotide-gated ion channel 6--is involved in heat shock responses. Plant J 70:1056–1069. 10.1111/j.1365-313X.2012.04969.x.

Gehring, C., and Turek, I.S. (2017). Cyclic Nucleotide Monophosphates and Their Cyclases in Plant Signaling. Front Plant Sci 8:1704. 10.3389/fpls.2017.01704.

Genschik, P., Hall, J., and Filipowicz, W. (1997). Cloning and characterization of the Arabidopsis cyclic phosphodiesterase which hydrolyzes ADP-ribose 1’’,2’’-cyclic phosphate and nucleoside 2’,3’-cyclic phosphates. Journal of Biological Chemistry 272:13211–13219. DOI 10.1074/jbc.272.20.13211.

Gerits, N., Mikalsen, T., Kostenko, S., Shiryaev, A., Johannessen, M., and Moens, U. (2007). Modulation of F-actin rearrangement by the cyclic AMP/cAMP-dependent protein kinase (PKA) pathway is mediated by MAPK-activated protein kinase 5 and requires PKA-induced nuclear export of MK5. J Biol Chem 282:37232–37243. 10.1074/jbc.M704873200.

Gilliland, L.U., Pawloski, L.C., Kandasamy, M.K., and Meagher, R.B. (2003). Arabidopsis actin gene ACT7 plays an essential role in germination and root growth. Plant J 33:319–328. 10.1046/j.1365-313x.2003.01626.x.

Henty-Ridilla, J.L., Shimono, M., Li, J., Chang, J.H., Day, B., and Staiger, C.J. (2013). The plant actin cytoskeleton responds to signals from microbe-associated molecular patterns. PLoS Pathog 9:e1003290. 10.1371/journal.ppat.1003290.

Horwich, A.L., Fenton, W.A., Chapman, E., and Farr, G.W. (2007). Two families of chaperonin: physiology and mechanism. Annu Rev Cell Dev Biol 23:115–145. 10.1146/annurev.cellbio.23.090506.123555.

Howe, A.K. (2004). Regulation of actin-based cell migration by cAMP/PKA. Biochim Biophys Acta 1692:159–174. 10.1016/j.bbamcr.2004.03.005.

Huang, S., Blanchoin, L., Kovar, D.R., and Staiger, C.J. (2003). Arabidopsis capping protein (AtCP) is a heterodimer that regulates assembly at the barbed ends of actin filaments. J Biol Chem 278:44832–44842. 10.1074/jbc.M306670200.

Jackson, E.K. (1991). Adenosine: a physiological brake on renin release. Annu Rev Pharmacol Toxicol 31:1–35. 10.1146/annurev.pa.31.040191.000245.

Jarratt-Barnham, E., Wang, L., Ning, Y., and Davies, J.M. (2021). The Complex Story of Plant Cyclic Nucleotide-Gated Channels. Int J Mol Sci 2210.3390/ijms22020874.

Jiang, J., Fan, L.W., and Wu, W.H. (2005). Evidences for involvement of endogenous cAMP in Arabidopsis defense responses to Verticillium toxins. Cell Res 15:585–592. 10.1038/sj.cr.7290328.

Kandasamy, M.K., McKinney, E.C., and Meagher, R.B. (2009). A single vegetative actin isovariant overexpressed under the control of multiple regulatory sequences is sufficient for normal Arabidopsis development. Plant Cell 21:701–718. 10.1105/tpc.108.061960.

Kandasamy, M.K., McKinney, E.C., Roy, E., and Meagher, R.B. (2012). Plant vegetative and animal cytoplasmic actins share functional competence for spatial development with protists. Plant Cell 24:2041–2057. 10.1105/tpc.111.095281.

Ketelaar, T., Allwood, E.G., Anthony, R., Voigt, B., Menzel, D., and Hussey, P.J. (2004). The actin-interacting protein AIP1 is essential for actin organization and plant development. Curr Biol 14:145–149. 10.1016/j.cub.2004.01.004.

Lefkimmiatis, K., Moyer, M.P., Curci, S., and Hofer, A.M. (2009). “cAMP sponge”: a buffer for cyclic adenosine 3’, 5’-monophosphate. PLoS One 4:e7649. 10.1371/journal.pone.0007649.

Leng, Q., Mercier, R.W., Hua, B.G., Fromm, H., and Berkowitz, G.A. (2002). Electrophysiological analysis of cloned cyclic nucleotide-gated ion channels. Plant Physiol 128:400–410. 10.1104/pp.010832.

Li, J., Wang, X., Qin, T., Zhang, Y., Liu, X., Sun, J., Zhou, Y., Zhu, L., Zhang, Z., Yuan, M., et al. (2011). MDP25, a novel calcium regulatory protein, mediates hypocotyl cell elongation by destabilizing cortical microtubules in Arabidopsis. Plant Cell 23:4411–4427. 10.1105/tpc.111.092684.

Li, W., Luan, S., Schreiber, S.L., and Assmann, S.M. (1994). Cyclic AMP stimulates K+ channel activity in mesophyll cells of Vicia faba L. Plant Physiol 106:957–961. 10.1104/pp.106.3.957.

Lim, Y.T., Prabhu, N., Dai, L., Go, K.D., Chen, D., Sreekumar, L., Egeblad, L., Eriksson, S., Chen, L., Veerappan, S., et al. (2018). An efficient proteome-wide strategy for discovery and characterization of cellular nucleotide-protein interactions. PLoS One 13:e0208273. 10.1371/journal.pone.0208273.

Lu, M., Zhang, Y., Tang, S., Pan, J., Yu, Y., Han, J., Li, Y., Du, X., Nan, Z., and Sun, Q. (2016). AtCNGC2 is involved in jasmonic acid-induced calcium mobilization. J Exp Bot 67:809–819. 10.1093/jxb/erv500.

Ma, W., and Berkowitz, G.A. (2011). Ca2+ conduction by plant cyclic nucleotide gated channels and associated signaling components in pathogen defense signal transduction cascades. New Phytol 190:566–572. 10.1111/j.1469-8137.2010.03577.x.

Makman, R.S., and Sutherland, E.W. (1965). Adenosine 3’,5’-Phosphate in Escherichia Coli. J Biol Chem 240:1309–1314.

Mao, T., Jin, L., Li, H., Liu, B., and Yuan, M. (2005). Two microtubule-associated proteins of the Arabidopsis MAP65 family function differently on microtubules. Plant Physiol 138:654–662. 10.1104/pp.104.052456.

Moutinho, A., Hussey, P.J., Trewavas, A.J., and Malho, R. (2001). cAMP acts as a second messenger in pollen tube growth and reorientation. Proceedings of the National Academy of Sciences of the United States of America 98:10481–10486. 10.1073/pnas.171104598.

Perez-Riverol, Y., Csordas, A., Bai, J., Bernal-Llinares, M., Hewapathirana, S., Kundu, D.J., Inuganti, A., Griss, J., Mayer, G., Eisenacher, M., et al. (2019). The PRIDE database and related tools and resources in 2019: improving support for quantification data. Nucleic Acids Res 47:D442–D450. 10.1093/nar/gky1106.

Porter, K., and Day, B. (2016). From filaments to function: The role of the plant actin cytoskeleton in pathogen perception, signaling and immunity. J Integr Plant Biol 58:299–311. 10.1111/jipb.12445.

Qian, D., and Xiang, Y. (2019). Actin Cytoskeleton as Actor in Upstream and Downstream of Calcium Signaling in Plant Cells. Int J Mol Sci 2010.3390/ijms20061403.

Qin, T., Liu, X., Li, J., Sun, J., Song, L., and Mao, T. (2014). Arabidopsis microtubule-destabilizing protein 25 functions in pollen tube growth by severing actin filaments. Plant Cell 26:325–339. 10.1105/tpc.113.119768.

Rall, T.W., and Sutherland, E.W. (1958). Formation of a cyclic adenine ribonucleotide by tissue particles. J Biol Chem 232:1065–1076.

Rayapuram, N., Bigeard, J., Alhoraibi, H., Bonhomme, L., Hesse, A.M., Vinh, J., Hirt, H., and Pflieger, D. (2018). Quantitative Phosphoproteomic Analysis Reveals Shared and Specific Targets of Arabidopsis Mitogen-Activated Protein Kinases (MAPKs) MPK3, MPK4, and MPK6. Mol Cell Proteomics 17:61–80. 10.1074/mcp.RA117.000135.

Reiss, H.D., and Herth, W. (1985). Nifedipine-sensitive calcium channels are involved in polar growth of lily pollen tubes. J Cell Sci 76:247–254.

Sabetta, W., Vannini, C., Sgobba, A., Marsoni, M., Paradiso, A., Ortolani, F., Bracale, M., Viggiano, L., Blanco, E., and de Pinto, M.C. (2016). Cyclic AMP deficiency negatively affects cell growth and enhances stress-related responses in tobacco Bright Yellow-2 cells. Plant Mol Biol 90:467–483. 10.1007/s11103-016-0431-5.

Sabetta, W., Vandelle, E., Locato, V., Costa, A., Cimini, S., Bittencourt Moura, A., Luoni, L., Graf, A., Viggiano, L., De Gara, L., et al. (2019). Genetic buffering of cyclic AMP in Arabidopsis thaliana compromises the plant immune response triggered by an avirulent strain of Pseudomonas syringae pv. tomato. Plant J 98:590–606. 10.1111/tpj.14275.

Savitski, M.M., Reinhard, F.B., Franken, H., Werner, T., Savitski, M.F., Eberhard, D., Martinez Molina, D., Jafari, R., Dovega, R.B., Klaeger, S., et al. (2014). Tracking cancer drugs in living cells by thermal profiling of the proteome. Science 346:1255784. 10.1126/science.1255784.

Shannon, P., Markiel, A., Ozier, O., Baliga, N.S., Wang, J.T., Ramage, D., Amin, N., Schwikowski, B., and Ideker, T. (2003). Cytoscape: a software environment for integrated models of biomolecular interaction networks. Genome Res 13:2498–2504. 10.1101/gr.1239303.

Smith, C.J., Brown, E.G., Newton, R.P., Al-Najafi, T., and Edwards, M.J. (1977). Adenylate cyclase activity in higher plants [proceedings]. Biochem Soc Trans 5:1351–1353. 10.1042/bst0051351.

Sy, J., and Richter, D. (1972). Separation of a cyclic 3’,5’-adenosine monophosphate binding protein from yeast. Biochemistry 11:2784–2787. 10.1021/bi00765a008.

Talke, I.N., Blaudez, D., Maathuis, F.J., and Sanders, D. (2003). CNGCs: prime targets of plant cyclic nucleotide signalling? Trends Plant Sci 8:286–293. 10.1016/S1360-1385(03)00099-2.

Topf, A., Franz, P., and Tsiavaliaris, G. (2017). MicroScale Thermophoresis (MST) for studying actin polymerization kinetics. Biotechniques 63:187–190. 10.2144/000114599.

Totsuka, Y., Ferdows, M.S., Nielsen, T.B., and Field, J.B. (1983). Effects of forskolin on adenylate cyclase, cyclic AMP, protein kinase and intermediary metabolism of the thyroid gland. Biochim Biophys Acta 756:319–327. 10.1016/0304-4165(83)90340-9.

Turnham, R.E., and Scott, J.D. (2016). Protein kinase A catalytic subunit isoform PRKACA; History, function and physiology. Gene 577:101–108. 10.1016/j.gene.2015.11.052.

von Mering, C., Huynen, M., Jaeggi, D., Schmidt, S., Bork, P., and Snel, B. (2003). STRING: a database of predicted functional associations between proteins. Nucleic Acids Res 31:258–261.

Walter, L.M., Franz, P., Lindner, R., Tsiavaliaris, G., Hensel, N., and Claus, P. (2020). Profilin2a-phosphorylation as a regulatory mechanism for actin dynamics. FASEB J 34:2147–2160. 10.1096/fj.201901883R.

Wang, Y., Zhu, Y., Ling, Y., Zhang, H., Liu, P., Baluska, F., Samaj, J., Lin, J., and Wang, Q. (2010). Disruption of actin filaments induces mitochondrial Ca2+ release to the cytoplasm and [Ca2+]c changes in Arabidopsis root hairs. BMC Plant Biol 10:53. 10.1186/1471-2229-10-53.

Wang, Y.F., Fan, L.M., Zhang, W.Z., Zhang, W., and Wu, W.H. (2004). Ca2+-permeable channels in the plasma membrane of Arabidopsis pollen are regulated by actin microfilaments. Plant Physiol 136:3892–3904. 10.1104/pp.104.042754.

Xu, X.M., Wang, J., Xuan, Z., Goldshmidt, A., Borrill, P.G., Hariharan, N., Kim, J.Y., and Jackson, D. (2011). Chaperonins facilitate KNOTTED1 cell-to-cell trafficking and stem cell function. Science 333:1141–1144. 10.1126/science.1205727.

Yau, K.W. (1994). Cyclic nucleotide-gated channels: an expanding new family of ion channels. Proc Natl Acad Sci U S A 91:3481–3483. 10.1073/pnas.91.9.3481.

Yoshioka, K., Moeder, W., Kang, H.G., Kachroo, P., Masmoudi, K., Berkowitz, G., and Klessig, D.F. (2006). The chimeric Arabidopsis CYCLIC NUCLEOTIDE-GATED ION CHANNEL11/12 activates multiple pathogen resistance responses. Plant Cell 18:747–763. 10.1105/tpc.105.038786.

Zelman, A.K., Dawe, A., Gehring, C., and Berkowitz, G.A. (2012). Evolutionary and structural perspectives of plant cyclic nucleotide-gated cation channels. Front Plant Sci 3:95. 10.3389/fpls.2012.00095.

